# Wheat *EARLY FLOWERING3* is a dawn-expressed circadian oscillator component that regulates heading date

**DOI:** 10.1101/2021.09.03.458922

**Authors:** Lukas Wittern, Gareth Steed, Laura J. Taylor, Dora Cano Ramirez, Gabriela Pingarron-Cardenas, Keith Gardner, Andy Greenland, Matthew A. Hannah, Alex A. R. Webb

**Affiliations:** Department of Plant Sciences, University of Cambridge, Downing Street, Cambridge, CB2 3EA, UK; NIAB, Huntingdon Road, Cambridge, UK; BASF, BBCC – Innovation Center Gent, Technologiepark-Zwijnaarde, Gent, Belgium

## Abstract

Using an eight-parent Multiparent Advanced Generation Inter-Cross (MAGIC) population we investigated how variation at circadian clock-associated genes contributes to the regulation of heading date in UK and European winter wheat varieties. We identified homoeologues of *EARLY FLOWERING* 3 (*ELF3*) as candidates for the *Earliness per se* (*Eps*) *D1* and *B1 loci* in field conditions. We confirmed that a SNP within the coding region of *TaELF3-B1* is a candidate polymorphism underlying the *Eps-B1 locus.* We found that a reported deletion at the *Eps-D1 locus* encompassing *TaELF3-D1,* is instead a novel allele that lies within an introgression region containing an inversion relative to the Chinese Spring D genome. Using *T. turgidum cv. Kronos* carrying loss of function alleles of *TtELF3* we show that *ELF3* does regulate heading by demonstrating that the loss of a single *ELF3* homoeologue was sufficient to alter heading date. These studies demonstrated that *ELF3* forms part of the circadian oscillator but loss of all homoeologues was required to affect circadian rhythms. Similarly, loss of functional *LUX ARRHYTHMO* (*LUX*) in *T. aestivum*, an orthologue of a protein partner of Arabidopsis ELF3, severely disrupted circadian rhythms. *ELF3* and *LUX* transcripts are not co-expressed at dusk suggesting the structure of the wheat circadian oscillator might differ to that of Arabidopsis. Our demonstration that alteration to *ELF3* homoeologues can affect heading date separate from effects on the circadian oscillator suggests a role for *ELF3* in cereal photoperiodic responses that could be selected for, without pleiotropic deleterious alterations to circadian rhythms.

## Introduction

Circadian clocks are endogenous timing mechanisms with a near 24 hour period that sequence biological events, provide anticipation of environmental rhythms and permit measurement of daylength to regulate seasonal adjustments (McClung 2019). Plant breeders have indirectly selected for variation at circadian-associated *loci* in many of the world’s major crops when breeding to increase yield and improve crop performance by adapting genotypes to the local environment (Steed et al. 2021). Here we investigated whether variation at circadian-associated *loci* contributes to wheat yield traits in the field using a Multiparent Advanced Generation Inter-Cross (MAGIC) mapping population (Mackay *et al*. 2014). Wheat is a major crop, providing approximately 20% of the dietary calories and protein for the world’s population (Hawkesford *et al*. 2013). Since the green revolution wheat yields have continued to increase but with the world’s population estimated to reach 9.6 billion in 2050 (Gerland *et al*. 2014) it is likely that demand will outpace predicted yield increases (Hawkesford *et al*. 2013). Identification of novel genetic variation contributing to yield is critical to increasing the rate of yield gains in wheat.

Understanding of the nature of plant circadian oscillators is based primarily on investigation of Arabidopsis. In that model, the circadian oscillator contains a series of interlocking transcriptional feedback loops. The circadian oscillator cycle can be considered to start at dawn when expression of the transcriptional repressors *CIRCADIAN CLOCK ASSOCIATED 1* (*CCA1*) and *LATE ELONGATED HYPOCOTYL* (*LHY*) peaks, followed sequentially by peaks of expression of *PSEUDO RESPONSE REGULATOR 9* (*PRR9*)*, PRR7, PRR5, PRR3* and finally *PRR1* (also known as*TIMING OF CAB EXPRESSION 1* (*TOC1*)) at dusk (Hsu and Harmer 2014). PRR9 and PRR7 repress the expression of *CCA1* and *LHY* which releases repression of evening phased genes including, *TOC1, GIGANTEA* (*GI*)*, EARLY FLOWERING 3* (*ELF3*)*, ELF4* and *LUX ARRHYTHMO* (*LUX*) (Hsu and Harmer 2014). LUX is a MYB-like transcription factor which associates with ELF3 and ELF4 to form the tripartite evening complex (EC), which represses expression of *GI, TOC1, PRR7, PRR9* and *LUX* (Dixon *et al*. 2011, Nusinow *et al*. 2011, Herrero *et al*. 2012). Mutation of any of the three EC components causes arrhythmia of the Arabidopsis circadian oscillator in continuous light and temperature (Nagel and Kay 2012).

Orthologues of *ELF3* underlie *loci* regulating heading date in multiple crops, including barley *early maturity8* (*eam8*) (Faure *et al*. 2012), rice *heading date17* (*Hd17*) (Matsubara *et al*. 2012), *Earliness per se* (*Eps*)-*A^m^1* in *T. monococcum* and *Eps-D1* in *T. aestivum* (Zikhali *et al*. 2016). In hexaploid wheat, the magnitude of the *Eps-D1* effect on heading date increases with decreasing temperature (Ochagavía *et al*. 2019). Orthologues of *TOC1* are associated with heading date, plant height and thousand grain weight (Sun *et al*. 2020). An orthologue of *LUX* is a candidate for *Eps-3A^m^* in *T. monococcum* (Gawronski *et al*. 2014) and *HvLUX* is a candidate gene for *eam10* in barley (Campoli *et al*. 2013). Furthermore, the *Photoperiod-1* (*Ppd-1*) genes, which contribute to photoperiodic regulation of ear emergence, are orthologues of Arabidopsis *PRRs* (Beales *et al*. 2007, Shaw *et al*. 2012, Bentley *et al*. 2013, Turner *et al*. 2013).

We used the allelic diversity of a wheat MAGIC population to investigate the role of circadian clock genes in regulating yield component traits in the field. The parent varieties for the MAGIC population used in this study (Alchemy, Brompton, Claire, Hereward, Rialto, Robigus, Soissons and Xi-19) are representative of UK and European winter wheat varieties, capturing >80% of the polymorphism found across UK wheat varieties from the last 70 years (Gardner *et al*. 2016). Our data demonstrate that modified *ELF3* function can affect the regulation of flowering time without disrupting circadian oscillations and thus potentially avoid associated yield penalties (Dodd *et al*. 2005). Together, this suggests *ELF3* is a possible breeding target for crop improvement. We find evidence that in wheat *ELF3* is expressed in the morning, rather than the evening as it is in Arabidopsis and conclude there are likely differences in the role for *ELF3* in the circadian system between wheat and Arabidopsis. Our finding of separable roles for *ELF3* in the circadian oscillator and the regulation of heading date suggest a direct role for ELF3 in photoperiodic regulation in cereals.

## Results

### Earliness per se *loci Eps-B1* and *Eps-D1* segregate in an eight-parent wheat MAGIC population

We performed QTL analysis on a set of replicated trial data for 784 recombinant inbred lines (RILs) for the NIAB Elite MAGIC population from the 2012/2013 and 2013/2014 growing seasons at the NIAB experimental farms in Cambridge, UK. Growth stage 55 (GS55, equivalent to heading date) (Zadoks *et al*. 1974) was chosen for analysis and QTL mapping to enable comparisons with previous studies (Griffiths *et al*. 2009). In both 2013 and 2014, Soissons, which contains the *Ppd-D1a* (photoperiod insensitive) allele, reached GS55 6-8 days earlier than any of the other MAGIC parents, which contain the *Ppd-D1b* allele (photoperiod sensitive) (Figure 1). Heading date of the other MAGIC parents varied over 5.5 days in 2013 and 7 days in 2014.

**Figure 1.**
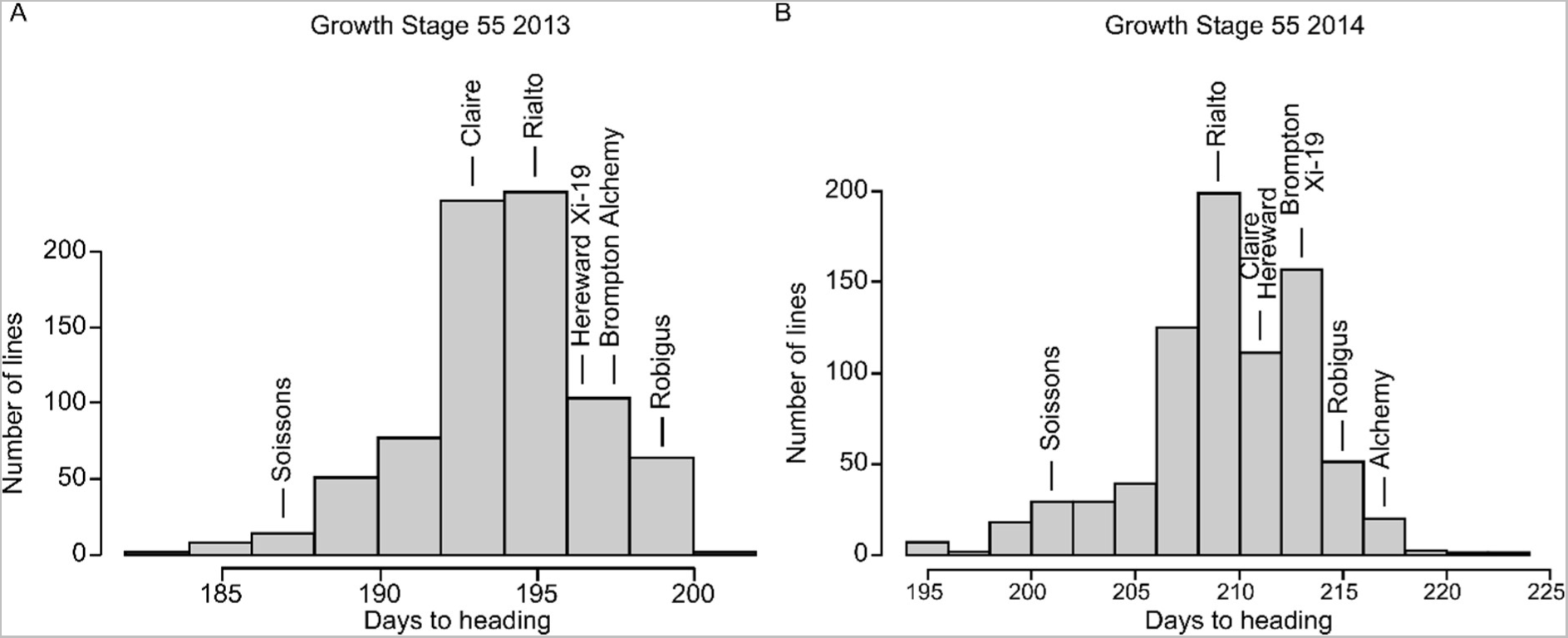
Variation in time to reach GS55 across a UK wheat MAGIC population. Time in days for individual MAGIC RILs (recombinant inbred lines) to reach GS55 for the (A) 2012/2013 growing season and (B) 2013/2014 growing season. The number of days after sowing to reach GS55 for the eight MAGIC parents is indicated by arrows.

We performed the QTL analysis using MPWGAIM (Verbyla *et al*. 2014) for GS55 phenotypes in both years and tested whether any circadian clock genes co-locate with the identified QTLs (< 2 cM to nearest 90 k marker or < 5 Mb to gene). Because we are interested in the effects of the circadian clock on yield traits, we report only QTLs associated with circadian clock genes. We found that four circadian gene orthologues (*TaELF3-B1, TaELF3-D1, Ppd-1, TaZTL-7A*) fell within 0.20, 0.27, 0.58 and 4.31 Mb respectively of one of the QTL flanking markers (Table 1). The absence of significant heading date QTLs linked to vernalisation genes such as *VRN1* confirmed that complete vernalisation had occurred, ensuring that observed variations in heading date were true *Earliness per-se* (*Eps*) effects rather than a variation in the vernalisation requirements between the MAGIC parents (Bentley *et al*. 2013, Camargo *et al*. 2016). A previous study identified a single GS55 QTL in the NIAB eight-parent MAGIC population (Camargo *et al*. 2016) but this difference in findings is likely a result of different growth conditions (greenhouse compared to field), lower replicate number (208 RILs in pots compared to 784 RILs in field plots) and the use of a different mapping methodology. Further validation of our analysis pipeline and approaches are described in the methods and reported in (Supplemental Table S1).

**Table 1:** Summary of circadian clock associated QTLs and their mapping positions. Output from univariate QTL model for the NIAB 2013 and 2014 GS55 data. Gene (Mb) refers to the BLAST hit starting coordinate of the gene’s cDNA against the IWGSCRefSeqv1.1. Dist (cM) refers to the markers genetic mapping distance in centiMorgan. Dist (Mb) refers to the BLAST hit starting coordinate of the marker sequence against the IWGSCRefSeqv1.1 LOGP is −log10(p) which corresponds to the overall significance of the QTLs as a measure of its strength. % var is the percentage of genetic variance contributed by QTL. Further description of methodology and column headers can be found in (Verbyla et al. 2014).

| Circadian clock gene | Gene (Mb) | Chr | L. Marker | dist (cM) | dist (Mb) | R. Marker | dist (cM) | dist (Mb) | Year | LOGP | % var |
| --- | --- | --- | --- | --- | --- | --- | --- | --- | --- | --- | --- |
| **TaELF3** | 685,646,575 | 1B | Ra_c109187_371 | 296.34 | 685,741,261 | RAC875_c37075_283 | 297 | 687,715,336 | 2013 | 4.48 | 3.4 |
|  |  | 1B | Ra_c109187_371 | 296.34 | 685,741,261 | RAC875_c37075_284 | 297 | 687,715,336 | 2014 | 3.2 | 2 |
| **TaELF3** | 493,485,612 | 1D | Kukri_c29687_369 | 100.95 | 493,212,138 | Kukri_rep_c69829_307 | 101.95 | 488,577,079 | 2014 | 2.32 | 4.1 |
| **Ppd-1** | 33,952,591 | 2D | Kukri_c27309_590 | 48.57 | 31,806,310 | BS00064538_51 | 56.64 | 33,370,535 | 2013 | 20.04 | 17.2 |
|  |  | 2D | Kukri_c27309_590 | 48.57 | 31,806,310 | BS00064538_51 | 56.64 | 33,370,535 | 2014 | 43.92 | 30.2 |
| **TaZTL** | 625,638,942 | 7A | Kukri_c6676_172 | 207.62 | 610,934,248 | Ku_c2990_1939 | 208.12 | 621,322,194 | 2013 | 2.87 | 2.6 |

A QTL on the distal end of chromosome 1B, delineated by the markers Ra_c109187_371 and RAC875_c37075_283, mapped closely to the physical position of *TaELF3-B1* (TraesCS1B02G477400) (Table 1). The 1B locus also had a homoeologous QTL on chromosome 1D delineated by the markers Kukri_c29687_369 and Kukri_rep_c69829_307 identifying *TaELF3-D1* (TraesCS1D02G451200) as a candidate gene (Table 1, Figure 2A). These QTLs co-locate with the previously described heading date QTLs *Eps-B1* and *Eps-D1* (Griffiths *et al*. 2009). In our data, *Eps-B1* had a significant effect in both years, explaining 3.4% of genetic variance in GS55 in 2013 and 2% in 2014 (Table 1). *Eps-D1* was also detected in both years but only had an overall significant effect (−log10(p) = 2.32) in 2014, accounting for 4.1% of genetic variance in GS55 in that year (Table 1). The *Eps-B1* and *Eps-D1 loci* were chosen for further in-depth analysis because of their proximity to *ELF3* homoeologues.

**Figure 2.**
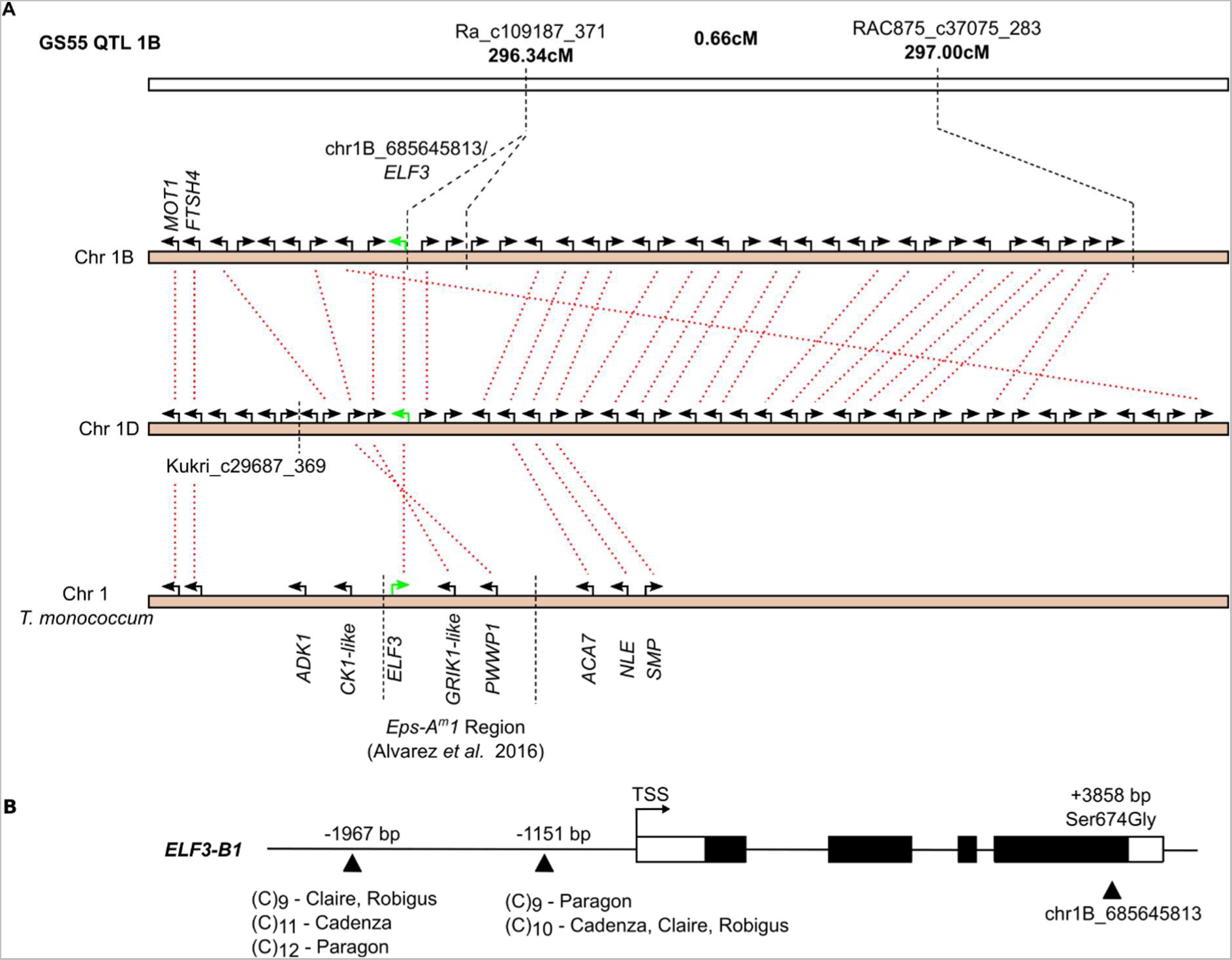
*ELF3* is a candidate gene for *Eps-B1*. (A) Syntenic relationships of the *Eps-B1* QTL to the *Eps-D1* QTL and the *T. monococcum Eps-A^m^1* QTL. *ELF3* highlighted in green. (B) *TaELF3-B1* gene model with exons represented by filled rectangles, UTR’s as white rectangles. The two promoter cytosine repeat polymorphisms with location relative to the transcription start site (TSS) are indicated with the cytosine (C) repeat number for Claire, Robigus, Cadenza and Paragon. The Ser674Gly SNP is situated within exon 4 of *ELF3-B1*.

### *TaELF3-B1* is a candidate gene for the *Eps-B1* QTL

At the 1B QTL, the alleles derived from the MAGIC parents Soissons, Alchemy, Robigus and Claire are associated with a relative reduction in the time to GS55, indicated by the negative sign of the QTL, whilst alleles from Rialto, Hereward, Xi-19 and Brompton are associated with a delay to GS55, indicated by the positive sign of the QTL (Table 2). The interval between the markers Ra_c109187_371 (296.34cM / 685.7Mb) and RAC875_c37075_283 (297.00cM / 687.7Mb) is 0.66cM, corresponding to a physical length of ca. 2 Mb (Figure 2A).

**Table 2.**
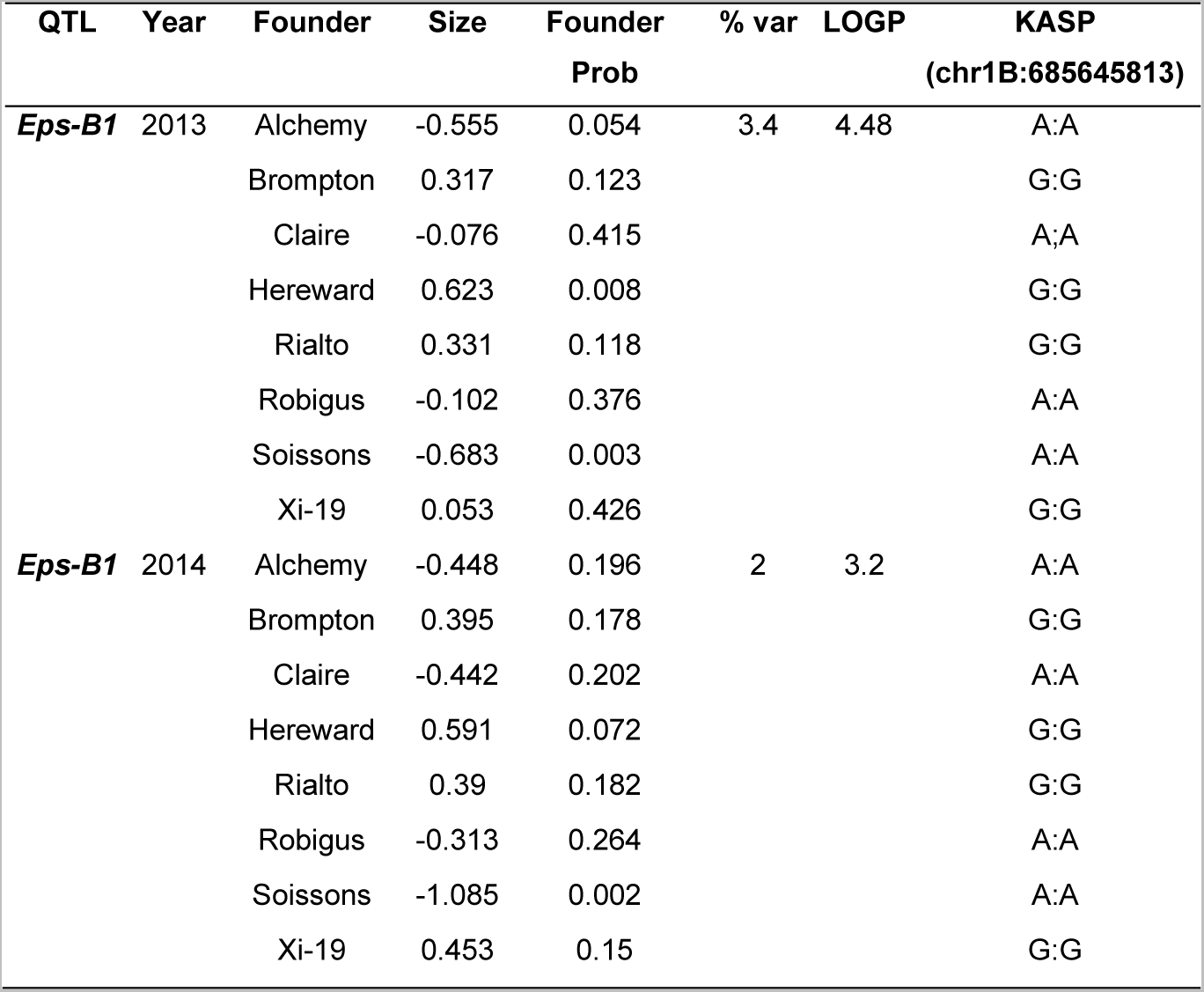
2013 and 2014 GS55 QTL summary for the *eps-B1 locus*. Estimated parental haplotype effects on RIL BLUPs. Abbreviations as per (Verbyla *et al*. 2014): Dist (cM) refers to the markers genetic mapping distance in centiMorgans (cM). LOGP is −log10(p) and corresponds to the overall significance of the QTL as a measure of its strength. Founder Prob: probability QTL is from founder shown (p). Strength of associations can thus be reported at the overall or founder level. Size: Size of effect of each founder allele (days to GS55).

| QTL | Year | Founder | Size | Founder Prob | % var | LOGP | KASP<br>(chr1B:685645813) |
| --- | --- | --- | --- | --- | --- | --- | --- |
| ***Eps-B1*** | 2013 | Alchemy | -0.555 | 0.054 | 3.4 | 4.48 | A:A |
|  |  | Brompton | 0.317 | 0.123 |  |  | G:G |
|  |  | Claire | -0.076 | 0.415 |  |  | A:A |
|  |  | Hereward | 0.623 | 0.008 |  |  | G:G |
|  |  | Rialto | 0.331 | 0.118 |  |  | G:G |
|  |  | Robigus | -0.102 | 0.376 |  |  | A:A |
|  |  | Soissons | -0.683 | 0.003 |  |  | A:A |
|  |  | Xi-19 | 0.053 | 0.426 |  |  | G:G |
| ***Eps-B1*** | 2014 | Alchemy | -0.448 | 0.196 | 2 | 3.2 | A:A |
|  |  | Brompton | 0.395 | 0.178 |  |  | G:G |
|  |  | Claire | -0.442 | 0.202 |  |  | A:A |
|  |  | Hereward | 0.591 | 0.072 |  |  | G:G |
|  |  | Rialto | 0.39 | 0.182 |  |  | G:G |
|  |  | Robigus | -0.313 | 0.264 |  |  | A:A |
|  |  | Soissons | -1.085 | 0.002 |  |  | A:A |
|  |  | Xi-19 | 0.453 | 0.15 |  |  | G:G |

**Table 3.** 2012 and 2014 GS55 QTL summary for the *eEps-D1 locus*. Estimated parental haplotype effects on RIL BLUPs. Abbreviations as per (Verbyla *et al*. 2014): Dist (cM) refers to the markers genetic mapping distance in centiMorgans (cM). LOGP is −log10(p) and corresponds to the overall significance of the QTL as a measure of its strength. Founder Prob: probability QTL is from founder shown (p). Strength of associations can thus be reported at the overall or founder level. Size: Size of effect of each founder allele (days to GS55).

| QTL | Year | Founder | Size | Founder Prob | % var | LOGP | PCR (Figureure 3D) |
| --- | --- | --- | --- | --- | --- | --- | --- |
| ***Eps-D1*** | 2012 | Alchemy | 0.552 | 0.254 | 0.9 | 0.56 | 1097bp |
|  |  | Brompton | 0.105 | 0.446 |  |  | 1097bp |
|  |  | Claire | -0.11 | 0.447 |  |  | 1097bp |
|  |  | Hereward | 0.021 | 0.49 |  |  | 1097bp |
|  |  | Rialto | -0.044 | 0.477 |  |  | 1097bp |
|  |  | Robigus | 0.003 | 0.498 |  |  | 1097bp |
|  |  | Soissons | 0.734 | 0.134 |  |  | 1097bp |
|  |  | Xi-19 | -1.261 | 0.015 |  |  | 776bp |
| ***Eps-D1*** | 2014 | Alchemy | -0.058 | 0.472 | 4.1 | 2.32 | 1097bp |
|  |  | Brompton | 0.869 | 0.104 |  |  | 1097bp |
|  |  | Claire | -1.412 | 0.051 |  |  | 1097bp |
|  |  | Hereward | -0.582 | 0.252 |  |  | 1097bp |
|  |  | Rialto | -0.071 | 0.457 |  |  | 1097bp |
|  |  | Robigus | -0.265 | 0.38 |  |  | 1097bp |
|  |  | Soissons | 1.244 | 0.013 |  |  | 1097bp |
|  |  | Xi-19 | -0.79 | 0.053 |  |  | 776bp |

The IWGSCv1.1 gene annotation suggests there are 26 high confidence gene models within the interval of Ra_c109187_371 and RAC875_c37075_283. Based on current gene annotation and orthology to cloned rice genes (Yao *et al*. 2018) there are no novel candidates for the *Eps-B1* locus in addition to those described previously (Faricelli *et al*. 2010, Alvarez *et al*. 2016, Zikhali *et al*. 2016). Compared to the *T. monococcum* physical map of the *Eps-A^m^1* region (Alvarez *et al*. 2016) *TmADK* (similar to rice putative kinase *ADK1)* and *TmCK1-like* (*CASEIN KINASE I ISOFORM DELTA-LIKE)* are not syntenic and the locus containing *TaELF3*, *TaGRIK1-like* (*GEMINIVIRUS REP INTERACTING KINASE 1 LIKE*) and *TaPWWP1* (*PWWP domain-containing protein*) is inverted (Figure 2A). Nineteen genes had homoeologues on the syntenic 1D chromosome (Figure 2A).

Publicly available sequences for *TaELF3-B1* in Claire, Robigus (earlier heading alleles) (Clavijo *et al*. 2017) and Rialto and Cadenza – the parents of Xi-19 - (later heading alleles) (Zikhali *et al*. 2016) differed by one nonsynonymous SNP within exon 4 at position chr1B:685645813. No further nonsynonymous SNPs were identified from the 820k SNP dataset from CerealsDB (Wilkinson *et al*. 2012), which contains all MAGIC parents.

The SNP (Adenine at position 2020 of CDS to Guanine; A2020G) causes a predicted deleterious amino acid substitution from the ancestral serine to a glycine residue at position 674 of the predicted primary amino acid sequence (Ser674Gly; SIFT 0.01) (Figure 2B). A KASP marker was designed for this SNP and used to genotype the MAGIC population. The KASP marker mapped to the same genetic position as Ra_c109187_371 and segregated with the predicted founder effects, confirming that *TaELF3-B1* is part of the 1B GS55 QTL.

This predicted glycine at residue 674 of *TaELF3-B1*, associated with late heading, is globally rare in wheat. In the 1000Exome data (He *et al*. 2019) 4.87% of lines have the predicted glycine residue and in the wheat HapMap panel (Jordan *et al*. 2015), only Rialto has the predicted glycine residue out of the 62 globally diverse lines analysed (Supplemental Table S2). Similarly, of the 14 hexaploid varieties currently available from the 10+ Wheat Genome project only three (Cadenza, Paragon, ArinaLrFor) possess the non-ancestral glycine residue (Supplemental Table S2). However, analysis of multiple datasets (see methods) found the non-ancestral glycine674 is more common in wheat varieties with a UK pedigree (9/13 varieties) (Supplemental Table S3). Alignment of *TaELF3* orthologues shows that the predicted serine residue is also highly conserved across monocots (Supplemental Figure S1).

When looking for other potentially candidate sequence variation we found that within the *ELF3-B1* promoter regions of Claire, Robigus, Paragon and Cadenza there are two sites polymorphic for the number of repeated cytosines ((C)_n_) located 1967 bp and 1151 bp upstream of the transcriptional start site (TSS) (Figure 2B). Variation in CT repeat number within the 5’-UTR region of the wheat *cellulose synthase like* (*Csl*) gene is associated with *Csl* expression and tiller number (Hyles *et al*. 2017). As Robigus and Claire have an early heading phenotype at the *Eps-B1* locus and Cadenza a late heading phenotype (Alvarez *et al*. 2016) it is unlikely that the promoter (C)_n_ polymorphisms cause the changes in heading dates because there is only a difference in two cytosines at the 1967 bp (C)_n_ site between Cadenza and Robigus/Claire (Figure 2B). This conclusion is supported by our finding that there is no correlation between the (C)_n_ genotype and *TaELF3* expression (ANOVA F = 0.484, p = 0.748) (Supplemental Figure S2). It was not possible to quantify the relative transcript abundance of the *TaELF3-B1* homoeologue due to a lack of homoeologous SNPs in the coding region of *TaELF3* as previously reported (Zikhali *et al*. 2016). Furthermore *TaELF3-B1* in Julius, Lancer, Mace, Norin61 and Sy Mattis seems to be inverted compared to ArinaLrFor, CDC Landmark, CDC Stanley, Chinese Spring and Jagger. However, as none of these lines share the same haplotype with ArinaLrFor and Paragon (which both carry the Ser674Gly SNP) at *ELF3-B1* it is unlikely that an inversion is the cause behind the *Eps-B1* QTL. Thus, we conclude that the Ser674Gly SNP in *ELF3-B1* is the best candidate to underlie *Eps-B1*.

### Subtelomeric introgression and inversion might underlie the *Eps-D1* heading date QTL

A subtelomeric deletion in Cadenza and Spark was previously described as the causal polymorphism underlying the *Eps-D1* locus using Spark × Rialto, Avalon × Cadenza mapping populations (Zikhali *et al*. 2016, Ochagavía *et al*. 2019). Our QTL analysis for 2012 (Supplemental Table S1) and 2014 (Table 1) shows a minor QTL overlapping this locus. In both years the Xi-19 founder genotype is associated with earlier heading (2012: −1.26 days; p = 0.015; 2014; −0.79 days; p = 0.053). However correct genetic map construction and resulting assignment of founder genotypes at this locus might be more difficult given the high-density marker region previously identified at the end of 1D. These high-density marker regions were hypothesised to be associated with introgressions (Gardner et al., 2016). Based on the previously reported deletion in Cadenza and Spark (Zikhali, Wingen, and Griffiths 2016) and the pedigree of Xi-19 (Cadenza/Rialto//Cadenza) we hypothesised that the Cadenza allele inherited from Tonic was a likely candidate for the observed QTL effects. By analysing data from recently published wheat assemblies (Walkowiak *et al*. 2020) we suggest the polymorphism is due to a subtelomeric inversion derived from introgressed material rather than a deletion. A BLAST search of the Chinese Spring *TaELF3-D1* genomic sequence against the 10+ Wheat Genome Project varieties identified high confidence matches in all varieties including Cadenza. Interestingly, the sequence retrieved from the variety Jagger was identical to Cadenza. However, the sequence identity of the Jagger and Cadenza coding sequence for TraesJAG1D01G492900.1 and TraesCAD_scaffold_026352_01G000400.1 compared to the Chinese Spring gene is much lower (96.72%) than for the remaining 1D coding sequences (>99.9%). The Jagger and Cadenza 1D coding sequences have a greater similarity to the Chinese Spring *TaELF3-A1* (97.50%) and Chinese Spring *TaELF3-B1* (97.88%) homoeologues than to Chinese Spring *TaELF3-D1*.

Visualisation of the 1D chromosome for the 10+ Wheat Genome Project varieties using the Crop Haplotypes viewer (Brinton *et al*. 2020) showed a shared haplotype block at the distal end of chromosome 1D between Cadenza (Haplotype viewer coordinates: 482.8 Mbp – 495.4 Mbp) and Jagger (480.0 Mbp to 493.5 Mbp) where *TaELF3*-*D1* is found.

The Cadenza v1.1 reference sequence has yet to be assembled into chromosomes, but since the Crop Haplotypes viewer (Brinton *et al*. 2020) data indicated identical by state (IBS) haplotypes between Jagger and Cadenza, we compared the distal end structure of the Jagger 1D chromosome with Chinese Spring. The end of the Jagger chromosome contains an inversion of approximately 2.6 Mbp relative to Chinese Spring which includes *TaELF3-D1* (Figure 3A). Alignment of the Jagger/Cadenza *TaELF3-D1* genomic sequence with Chinese Spring *TaELF3-D1* showed some sequence differences including a deletion within intron 2 (Figure 3B, Supplemental Table S4). We developed a subgenome specific PCR assay detailed in Figure 3B that amplifies the region of *TaELF3-D1* containing intron 2 and used this as a simple marker for the 1D introgression. We confirmed that of the eight MAGIC parents, only Xi-19 carried the Cadenza/Jagger *TaELF3-D1* allele (Figure 3C and D). As hypothesised Spark and Tonic also carry the Cadenza *TaELF3-D1* allele (Figure 3C, D and Supplemental Figure S3) as well as the varieties Maris Fundin and Cordiale which show introgression signatures on the distal end of chromosome 1D (Scott *et al*. 2021). A simplified pedigree diagram shows how some of the cultivars are related (Supplemental Figure S4). The intron 2 deletion is also present in *Aegilops speltoides* and *Triticum timopheevii* (Supplemental Figure S5).

**Figure 3.**
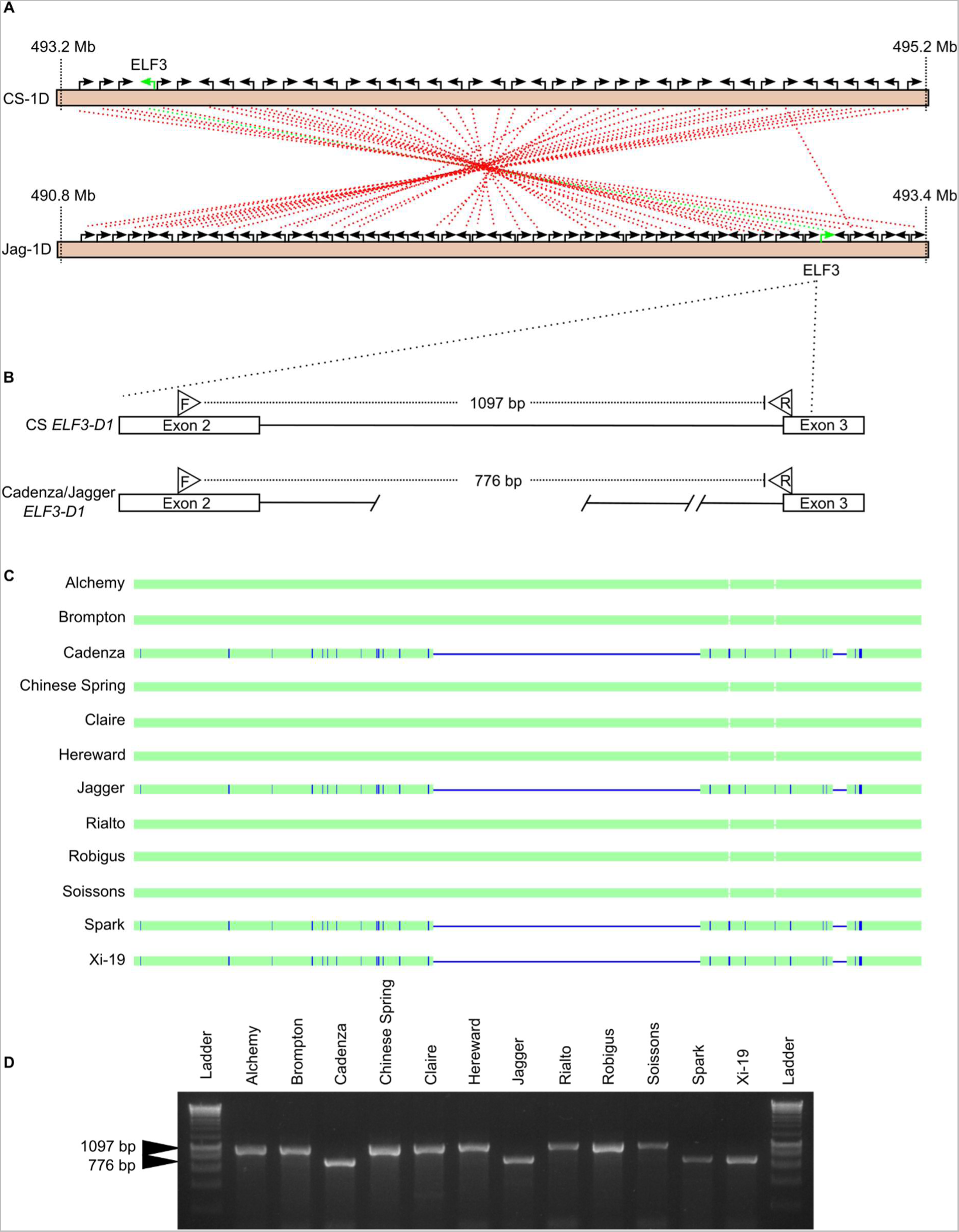
A subtelomeric chromosomal introgression and inversion containing *ELF3-D1* is likely to be the causal polymorphism underlying the *Eps-D1* QTL. (A) Syntenic relationships between the distal end of Chinese Spring (CS) 1D chromosome (IWGSCv1.1) and Jagger (Jag) 1D (PGSBv2.0), *TaELF3*-*D1* highlighted in green, each black arrow corresponds to agene in the forward (pointing right) or reverse (pointing left) strand. (B) the Cadenza/Jagger *ELF3-D1* contains a deletion within intron 2 which is not present in CS, F and R refer to relative primer binding sites for PCR amplification shown in (C). (D) Agarose gel separation of PCR products from (C). A PCR product at 776bp is indicative of the presence of a subtelomeric chromosomal introgression and inversion.

Given the low sequence conservation of the *TaELF3-D1* gene in Cadenza and Jagger and the presence of the intronic deletion in *Aegilops speltoides* and *Triticum timopheevii*, we investigated the wider phylogeny of the 1D Jagger/Cadenza region using the 820k SNP dataset from CerealsDB. This showed that Cadenza, Xi-19 and Spark as well as several other Tonic descendants (KWS-Podium, Cordiale, Gallant, Duxford) and five other cultivars (Moisson, Bacanora, Tuerkis, Badger, Pavon) clustered separately from the remaining *T. aestivum* cultivars (Supplemental Figure S6). These lines also cluster with six Watkins lines from India (Watkins34), Yugoslavia (Watkins352), China (Watkins141), UK (Watkins103), Cyprus (Watkins292) and Turkey (Watkins299). *T. timopheevii* clusters closely with the above lines. Gene alignment of *ELF3-D1* orthologues and alignment of 300 kb surrounding *ELF3-D1* also supports an origin outside the D genome for the Jagger/Cadenza *ELF3-D1* gene (Supplemental Figure S7, Supplemental Table S5).

### Allelic variation in *TaELF3* had no detectable effect on circadian rhythms in the MAGIC parent varieties

We investigated if the variations at *TaELF3-B1* and *TaELF3-D1* in the MAGIC parents, or other *loci,* might be associated with alterations of circadian rhythms in a manner that might explain the effects on heading date. We quantified circadian rhythms in the MAGIC parent varieties by measuring the period and amplitude of rhythms of delayed chlorophyll fluorescence (DF) in otherwise constant light (Gould *et al*. 2009). There were robust circadian rhythms of DF in all the MAGIC parent lines (RAE < 0.5) (Figure 4A). Hereward had the shortest circadian period (25.0 h) (Figure 4C) and Xi-19 the longest period (26.2 h) but the difference between these periods and the mean of all parents was not significant (p = 0.132) (Figure 4C). To determine whether the lack of an altered circadian phenotype in the MAGIC parent lines was the result of masking by compensatory alleles at other *loci,* we selected some of the earliest and latest heading MAGIC RILs from 2014 and used chlorophyll *a* fluorescence (PF) to measure circadian rhythms in continuous light (Supplemental Figure S8). There was no significant relationship between heading (early or late) and circadian period (p = 0.4216) (Supplemental Figure S8B) or between heading and circadian rhythm amplitude (p = 0.7385) (Supplemental Figure S8C). We detected robust rhythms of PF in all lines (RAE < 0.5) (Supplemental Figure S8D) indicating that in these lines there was no effect on circadian rhythm robustness. We then investigated whether there was any significant relationship between circadian rhythm parameters and the *ELF3-B1, ELF3-D1* or *Ppd-1* alleles associated with altered heading date. There was no significant difference between circadian period and *ELF3-B1* alleles (p = 0.542), *ELF3-D1* alleles (p = 0.521) or *Ppd-1* alleles (0.507). Similarly, there was no significant difference between circadian amplitude and *ELF3-B1* alleles (p = 0.942), *ELF3-D1* alleles (p = 0.1302) and *Ppd-1* alleles (0.966). Thus, the allelic diversity in homoeologues of *TaELF3* that was associated with alterations in heading date did not cause detectable changes in circadian rhythms.

**Figure 4.**
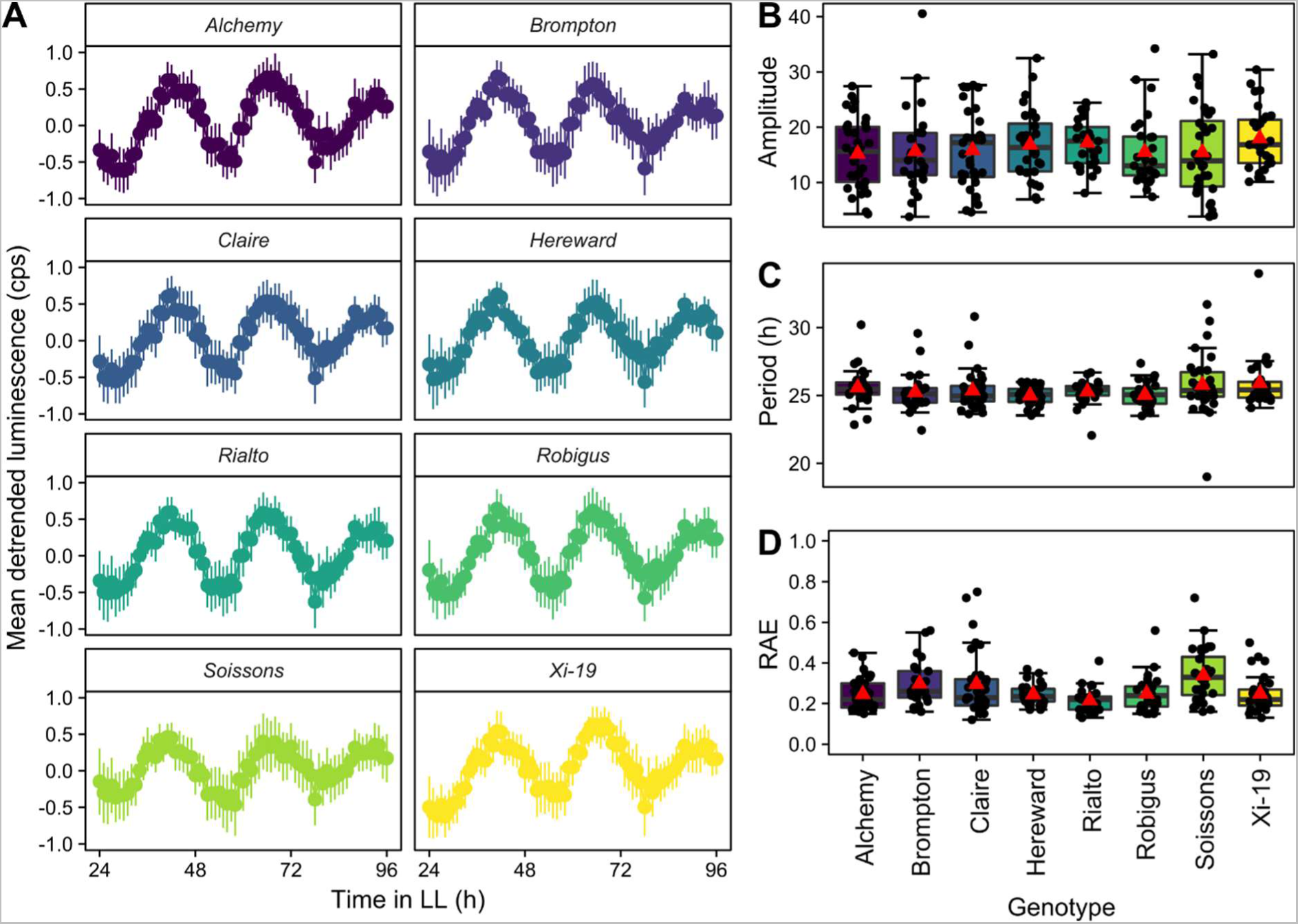
Circadian rhythms of delayed chlorophyll fluorescence in the MAGIC parents. (A) Mean DF luminescence in counts per second (cps) normalised to −1 to 1 using Biodare2 with error bars representing standard deviation (n = 27 - 33). Circadian amplitude (B), period (C) and RAE (D) calculated using FFT-NLLS (Biodare2). Upper and lower hinges represent the first and third quartiles (25^th^ and 75^th^ percentiles), the middle hinge represents the median value, red triangle represents meanvalue and black dots represent individual replicates.

### *TtELF3* has separable effects on heading date and the wheat circadian clock

Because allelic variation of *TaELF3* was associated with heading date but not circadian rhythms in the MAGIC parents and RILs, we examined directly whether *ELF3* affects heading and is a circadian clock component in wheat by examining tetraploid TILLING mutant lines carrying point mutations encoding stop codons in both *TtELF3-A1, TtELF3-B1* or both (Alvarez *et al*. 2016) (Supplemental Figure S9). The *Ttelf3*-null line headed first (48.7 days) with*Ttelf3*-WT heading significantly (p < 0.0001) later (56.4 days) as previously reported (Alvarez *et al*. 2016). There was no significant difference (p = 0.98) in heading date between *Ttelf3*-Anull (51.0 days) and *Ttelf3*-Bnull (52.4 days) suggesting that the *TtELF3-A* and *TtELF3-B* homoeologues contribute equally to heading date. However, they were both significantly later heading than *Ttelf3*-null (p < 0.0001) and significantly earlier than *Ttelf3*-WT (p < 0.0001).

We found that functional *ELF3* is required for robust circadian rhythms in continuous light and that single wild type *ELF3* homoeologues can maintain circadian function in the absence of other functional homoeologues. There were robust circadian rhythms of *F_m_’* in *Ttelf3*-WT (RAE 0.16 ± 0.05), *Ttelf3*-Anull (RAE 0.14 ± 0.04) and *Ttelf3*- Bnull (RAE 0.15 ± 0.07) but no robust circadian rhythms in *Ttelf3*-null (RAE 0.53 ± 0.19) (Figure 5B-C). The arrhythmic phenotype of *Ttelf3*-null was confirmed using DF and leaf temperature measurements (Supplemental Figure S10A-F) and was not the result of reduced maximum quantum yield of photosystem 2 (*F_v_/F_m_*) as this was higher for *Ttelf3*-null for the duration of measurement (Supplemental Figure S10G-I). There was no significant difference in circadian period of *F_m_’* (p > 0.05) between *Ttelf3*-WT (24.4 h ± 1.2), *Ttelf3*-Anull (24.3 h ± 0.7) and *Ttelf3*-Bnull (24.9 h ± 1.0). Similarly, there was no significant difference (p > 0.05) in the amplitude of *F_m_’* oscillations between *Ttelf3*-WT (241 ± 53), *Ttelf3*-Anull (218 ± 50) and *Ttelf3*-Bnull (236 ± 57).

**Figure 5.**
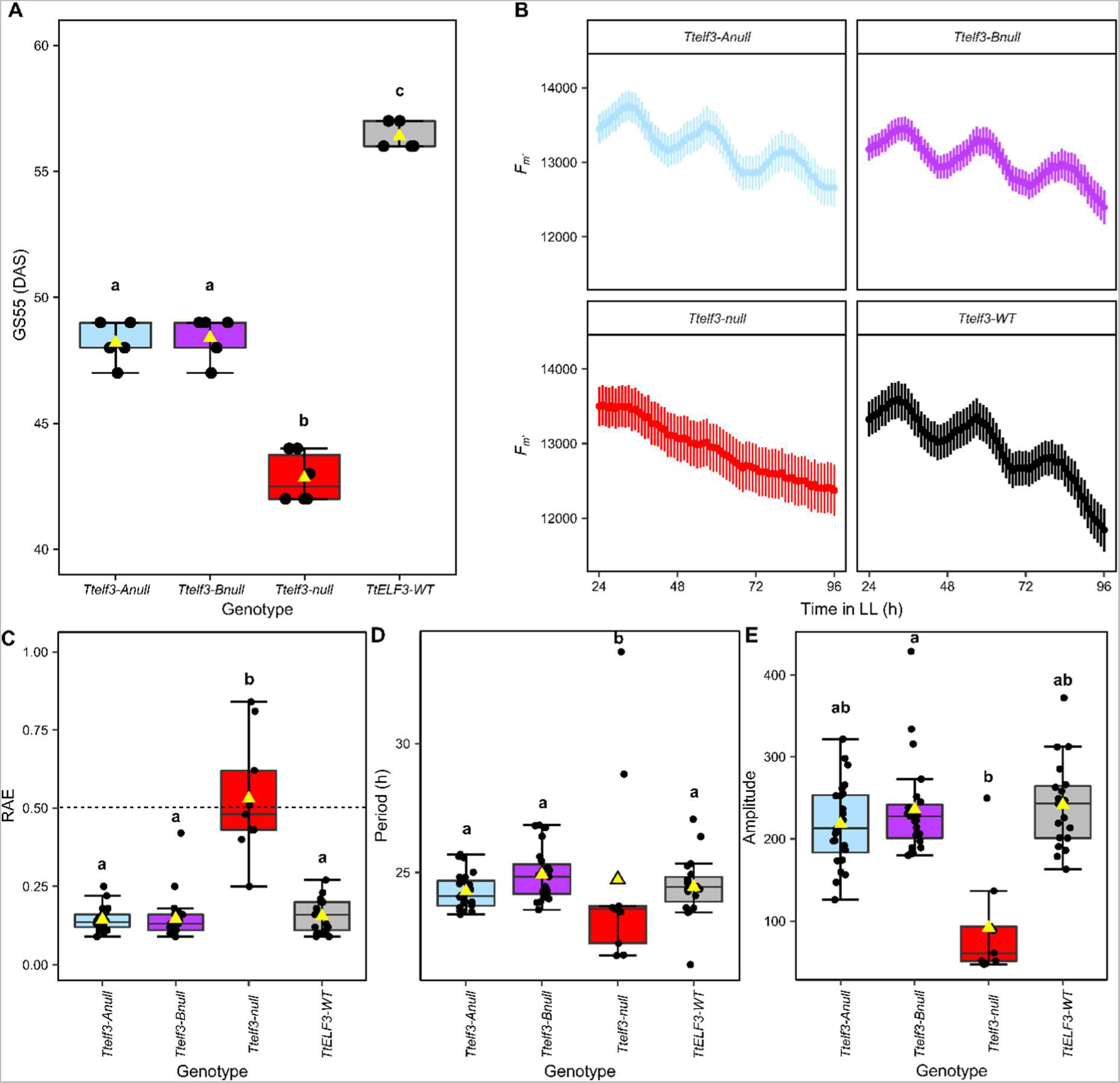
Mutation of single *ELF3* homoeologues affects heading date but not circadian rhythms of chlorophyll *a* fluorescence but *Ttelf3*-null is arrhythmic in continuous light. (A) GS55 for *Ttelf3*-null (red), *Ttelf3*-Anull (blue), *Ttelf3*-Bnull (purple) and *Ttelf3*-WT (grey) defined as the days after sowing (DAS) at which half of the first ear has emerged past the ligule. Upper and lower hinges represent the first and third quartiles (25^th^ and 75^th^ percentiles), the middle hinge represents the median value, yellow triangle represents mean value and black dots represent individual replicates (n = 5). (B) Mean *F_m_’* of lines shown in (A) in LL with SEM bars (n = 24). (C) RAE where points below dashed line considered rhythmic (D) circadian period length (hours)and (E) amplitude for genotypes inplot (B). Box jitter plots as described above, note plotted points in (C-E) correspond to those where data was successfully fitted by FFT-NLLS model in Biodare2. Significant differences (p < 0.05) calculated in R using Kruskal-Wallis test followed by post-hoc Dunn’s test.

Our results demonstrate that allelic variation at individual *ELF3* homoeologues is sufficient to alter heading date without affecting circadian rhythms, indicating that *ELF3* has separable effects on heading date and the circadian clock in wheat.

### Loss of functional *TtELF3* perturbs circadian oscillator gene expression in light/dark cycles and constant light

To further investigate the function of *ELF3* in the wheat circadian system we measured the relative transcript abundance of orthologues of the Arabidopsis circadian oscillator components *LHY, TOC1, Ppd-1* (*PRR37*)*, PRR73, GI, ELF3* and *LUX* in light dark (LD) and continuous light (LL) cycles. In LD cycles all transcripts had rhythmic expression (Figure 6A-G). Consistent with previous findings in Arabidopsis, *TtLHY* had peak expression at dawn (Figure 6A), the *PRR* orthologues peaked during the photoperiod (Figure 6B-C) with *TtTOC1* (Figure 6D), *TtGI* (Figure 6E) and *TtLUX* (Figure 6F) peaking around dusk. However, the peak in *TtELF3* abundance was at the end of the night just prior to dawn (Figure 6G), approximately antiphase to the expression of *TtLUX* (Figure 6F). This differs markedly from Arabidopsis in which *ELF3* and *LUX* are co-expressed near dusk (Nusinow *et al*. 2011, Huang *et al*. 2016, Ezer *et al*. 2017). For all transcripts, except *Ppd-1* (Figure 6B), we detected oscillations in relative transcript abundance in constant light, with the phasing in LL consistent with that in LD. These oscillations of circadian transcript abundance were abolished, or of very low amplitude, in *Ttelf3*-null (Figure 6A-G). Interestingly, the expression of *Ppd-1* was consistently higher in *Ttelf3*-null in both LD and the first LL cycle (Figure 6B) and the minimum level of the *Ppd-1* homologue *TtPRR73* was also higher in *Ttelf3*-null in LD (Figure 6C) which suggests that ELF3 might act as negative regulator of *Ppd-1* and *TtPRR73*.

**Figure 6.**
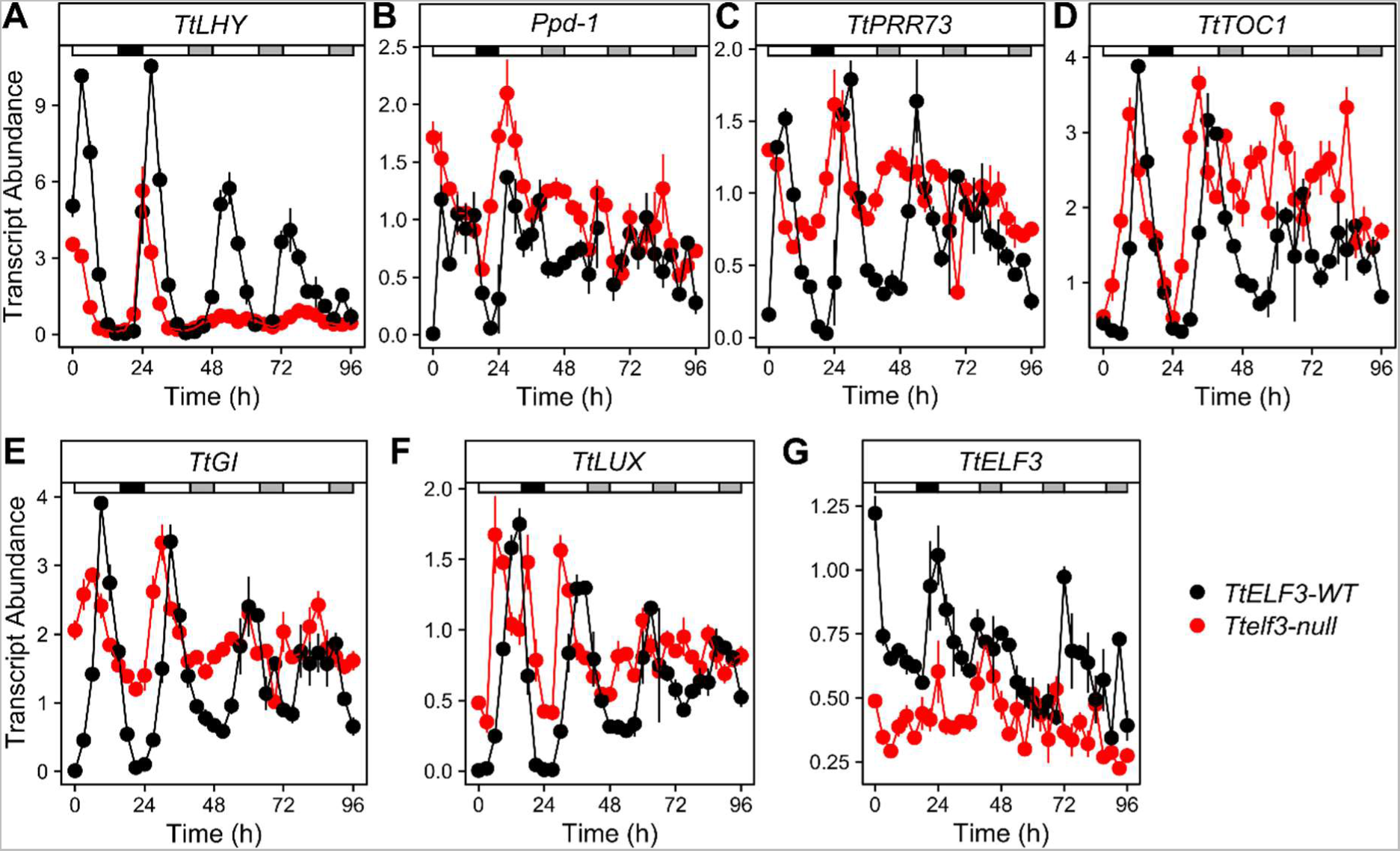
Abundance of wheat circadian clock transcripts in the *TtELF3-WT* and *ttelf3* lines in LD and LL. (A-G) Mean abundance of circadian oscillator transcripts (n= 3 - 5) in *Ttelf3*-WT (black) and *Ttelf3*- null (red) in a 24 h LD cycle in long day conditions (16 h light at 250 µmol m^−2^ s^−1^, 20°C: 8 h dark 16°C, represented by horizontal grey bar) followed by constant light and temperature (20°C) from time 24 to 96 h. Transcript abundance (ΔΔCq) is relative to *TtRP15* and *TtRPT5A* (A) *TtLHY,* (B) *Ppd-1,* (C) *TtPRR73,* (D) *TtTOC1,* (E) *TtGI,* (F) *TtLUX* and (G) *TtELF3*. White bar represents light, black bar represents darkness and grey bar represents subjective night.

There was a phase advance in peak transcript abundance of approximately 3 h in LD in *Ttelf3*-null compared to *TtELF3-WT* (Figure 6A-G) except for *TtELF3* transcript abundance where the peak phase was at dawn in both *Ttelf3*-WT and *Ttelf3*-null (Figure 6G). The amplitude of transcript abundance under diel conditions was also less in the *Ttelf3*-null line compared to *ELF3*-WT (Figure 6). Together, these data demonstrate that functional *TtELF3* is required for the maintenance of robust oscillations of circadian oscillator gene expression in LL and for normal circadian timing in LD.

### *LUX* loss of function disrupts the wheat circadian clock

The antiphase expression of wheat *ELF3* and *LUX* raises the possibility that the EC might not be formed, or functions differently to that in Arabidopsis. Previous studies in diploid wheat have identified *TmLUX* as a circadian oscillator gene (Gawronski *et al*. 2014), so we investigated whether *LUX* also contributes to circadian rhythms in hexaploid wheat, and whether functionality of one homoeologue is sufficient to maintain robust oscillations. We obtained three hexaploid Japanese cultivars, Chogokuwase, Minaminokomugi and Geurumil that have variation at *TaLUX* associated with flowering time (Mizuno *et al*. 2012). Chogokuwase is an extra early flowering variety and is the offspring of Geurumil and Minaminokomugi. Chogokuwase carries predicted non-functional mutations in all three sub-genome copies of *TaLUX* (Supplemental Figure S11). Chogokuwase also carries a *TaVRN-D1* spring allele and a photoperiod insensitive *Ppd-D1a* allele. Geurumil has the same predicted non-functional alleles of *TaLUX* and *Ppd-D1a* but has a winter growth habit due to lacking the *VRN-D1* spring allele. Minaminokomugi carries the non-functional *Talux-a1* allele but wild-type alleles of *TaLUX-B1* and *TaLUX-D1* in addition to carrying the *TaVRN-D1* and *Ppd-D1a* alleles (Mizuno *et al*. 2016).

Minaminokomugi maintained robust oscillations of chlorophyll fluorescence (RAE 0.16 ± 0.01) and leaf temperature (RAE 0.26 ± 0.02) (Figure 7A-C). In contrast Chogokuwase, which lacks functional copies of *TaLUX,* was arrhythmic for PF (RAE 0.56 ± 0.06) and Geurumil, which is also lacking functional *TaLUX*, had reduced robustness of PF oscillations (RAE 0.33 ± 0.05) compared to Minaminokomugi (Figure 7A-C). Similarly, there was a decreased robustness of oscillations in leaf surface temperature for Chogokuwase and Geurumil (RAE 0.48 ± 0.09 and RAE 0.46 ± 0.11) which appeared to dampen to arrhythmia after the first two true LL cycles (Figure 7D-F). We conclude that functional *TaLUX* is required for robust circadian rhythms in hexaploid wheat.

**Figure 7.**
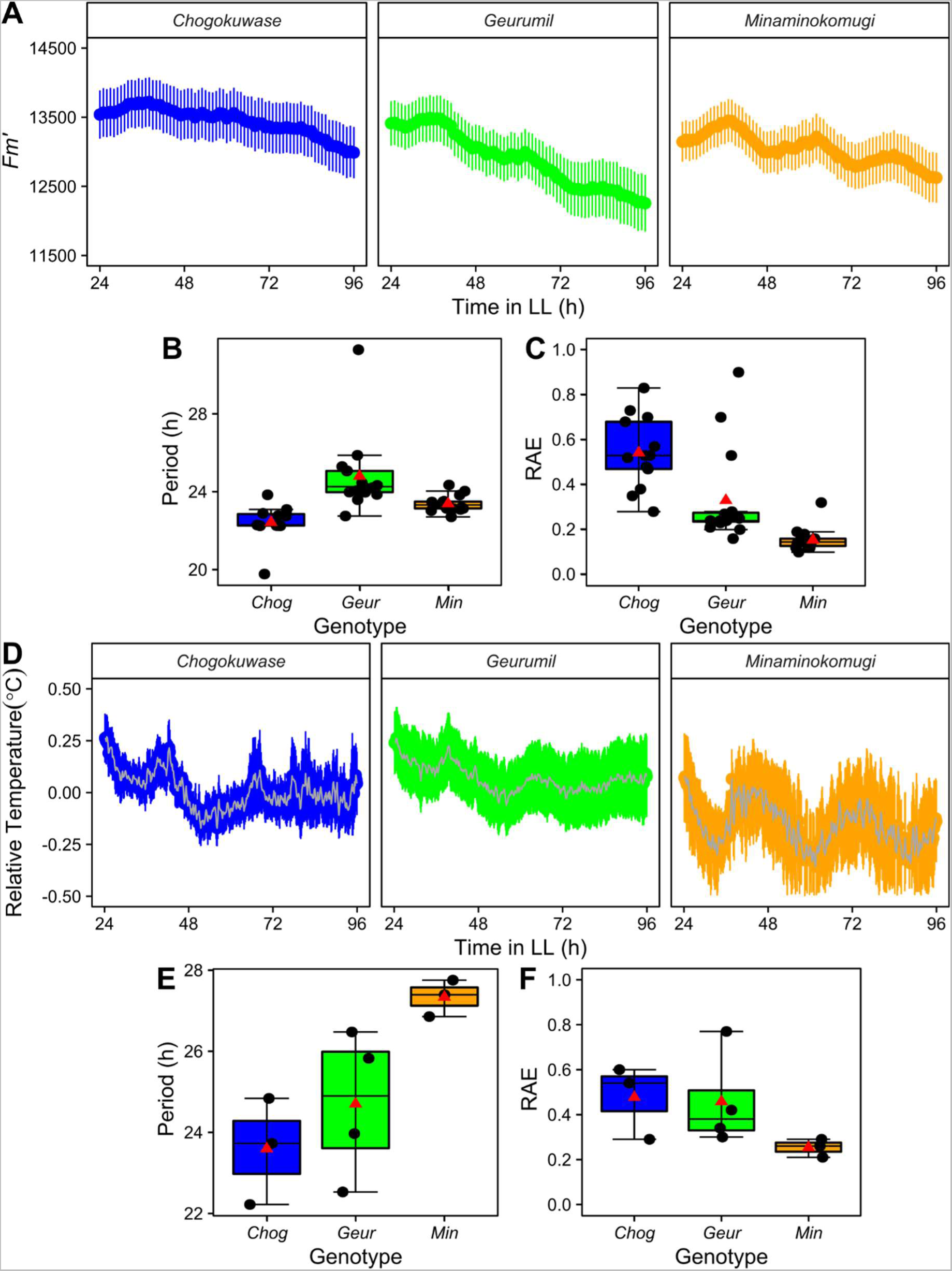
Functional *LUX* is required for maintenance of robust circadian oscillations under LL conditions in hexaploid wheat. (A) Mean *F_m_’* for Chogokuwase (blue), Geurumil (green) and Minaminokomugi (orange) in LL (n = 16)(B) Mean period and (C) RAE for (A) calculated using FFT-NLLS (Biodare2). Upper and lower hinges represent the first and third quartiles (25^th^ and 75^th^ percentiles), the middle hinge represents the median value, red triangle represents mean value and black dots represent individual replicates. (D) Mean Chogokuwase, Geurumil and Minaminokomugi leaf temperature relative to background in LL (n = 4).(E) mean period and (F) RAE for plot (D). Error bars are SEM, for (D) every 20^th^ point plotted for clarity.

## Discussion

### Wheat *ELF3* contributes to heading date and circadian rhythms independently

The eight MAGIC parent varieties provided the opportunity to measure the likely contribution of the *ELF3* homoeologues to heading date and circadian rhythms. Of the MAGIC parents Brompton, Hereward, Rialto and Xi-19 have a G|G substitution variant of *TaELF3-B1* and in addition Xi-19 has an inverted copy of *TaELF3-D1,* likely arising from a past introgression event. These variations in *TaELF3* alleles had no discernible impact on circadian rhythms (Figure 4) but were associated with measurable effects on heading date in the field (Supplemental Figure S12).

Consistent with previous studies, *ELF3* is a candidate for *Eps-D1 loci* (Zikhali *et al*. 2016). We have extended the findings concerning the *Eps-D1* locus by demonstrating that rather than being within a deletion, a copy of *ELF3-D1* is present in Cadenza (as well as Xi-19, Spark, Cordiale, Tonic and Maris Fundin) but does carry candidate sequence variations. *ELF3-D1* lies within an introgression region that contains an inversion relative to the Chinese Spring D genome (Figure 3) and current data cannot rule out other candidate genes within the introgression and/or inversion region. Based on analysis of the 820k SNP data we find that several descendants of the cultivar Tonic carry the putative introgression, as do several other unrelated cultivars and six Watkins lines from India, China, UK, Cyprus, Turkey, and Yugoslavia. The introgression is also likely present within Chinese elite germplasm (Wang *et al*. 2016). As a result, we hypothesise that this region predates modern breeding efforts. The same genomic region has been identified as being under selection in a collection of bread wheat accessions dating from 1790 to 1930, though in that study it was presumed to be a deletion rather than introgression (Przewieslik-Allen *et al*. 2021). While the origin of the introgression is uncertain, the presence of the Jagger/Cadenza *TaELF3-D1* intronic deletion in *Ae. speltoides* and *T. timopheevii* (Supplemental Figure S5) and the clustering outside of *Aegilops tauschii* lines (Supplemental Figure S6 and Supplemental Figure S7) supports an origin outside of the D genome, more closely related to the G or S genome. The increase in quantity and quality of genomic resources for diverse accessions and wheat relative species will help in determining the origin of this introgression.

Our data also identify *TaELF3-B1* as a candidate gene underlying the *Eps-B1* locus. We found that a predicted glycine674 (G|G) in *TaELF3-B1,* which segregates with *Eps-B1,* caused a delay in heading and flowering compared to the ancestral serine674 variant (A|A) (Supplemental Figure S12), in agreement with previous observations in Avalon (Zikhali *et al*. 2016). It was previously concluded that G|G is the ancestral variant (Zikhali *et al*. 2016) but our evidence suggests that it is more likely that A|A is ancestral because the G|G allele is globally rare, present in 4.87% of lines included in the 1000 Exome dataset, in one of the 62 wheat HapMap panel lines (Supplemental Table S2) and in three of the 14 10+ Wheat Genome project lines (Supplemental Table S2) and the ancestral serine residue is highly conserved, present in all sequenced grasses. However, within UK wheat germplasm (as defined by locality on wheat GRIS) the G|G allele is more common and is found in nine of the 13 UK lines we have investigated (Supplemental Table S3). The later flowering G|G allele may be more favoured in northern regions (Langer *et al*. 2014).

We found that the allelic variation at *TaELF3* had no detectable effect on circadian rhythms in the MAGIC parents or early and late heading RILs. Epistasis between *ELF3* and *Ppd-D1* has previously been reported for flowering time (Faure *et al*. 2012, Alvarez *et al*. 2016). However, we find that variation at *Ppd-A1* has no effects on circadian rhythms or modification of circadian phenotypes in *Ttelf3*-nulls (Supplemental Figure S13). We also find no evidence of epistasis between *eps-B1* and *eps-D1* in the regulation of circadian rhythms because Xi-19 has both *eps-B1* and *eps-D1* but was not distinguishable from the rest of the population in terms of circadian behaviour.

Using loss of function lines, we demonstrate that *ELF3* regulates heading and is required for circadian rhythms in wheat. Whilst complete loss of functional *TtELF3* caused arrhythmia of PF (Figure 5B-E & Supplemental Figure S10G-I), leaf temperature (Supplemental Figure S10D-F) and gene expression in LL (Figure 6), loss of function of either the A or B homoeologue was without effect on circadian rhythms if the other homoeologue was functional (Figure 5B-E). This contrasts with the effect on heading, where loss of either the A or B homoeologue function resulted in an advance in heading date relative to wildtype and the magnitude of this advance was intermediate relative to the double null mutant (Figure 5A). These results, coupled with the increased expression in *Ttelf3*-null of the floral promoter *Ppd-1* (Figure 6B) and *FT1* (Supplemental Figure S14), suggest that wheat *ELF3* functions as a core circadian oscillator component whilst also having a direct effect on flowering independent of its role in the circadian oscillator (Figure 8).

**Figure 8:**
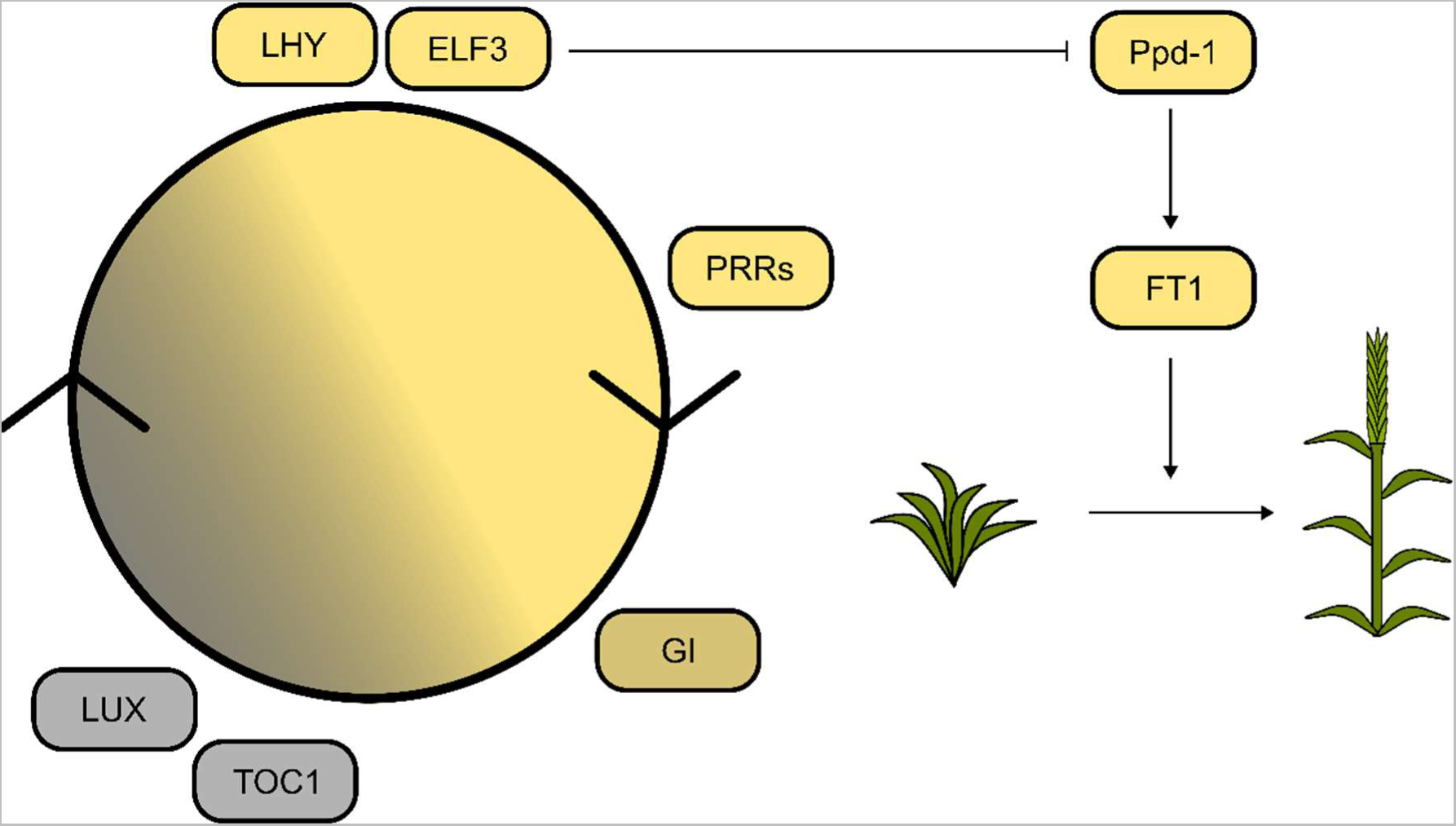
ELF3 functions in the circadian oscillator and regulates heading of wheat. Genes associated with the circadian oscillator are positioned and coloured relative to the time of day at which their transcript abundance is highest in LD. The progression of the circadian oscilaltor is indicated by the circle. Lines indicate potential regulatory links with outputs and other pathways. Pointed arrow heads represent activation and blunt arrowhead represents repression.

Loss of functional *TtELF3* advanced circadian phase in LD. This demonstrates that one role of *ELF3* is to contribute to the correct alignment of internal phase to external phase, which is a primary function of circadian oscillators. The earlier phasing of gene expression in LD cycles is similar to barley containing the *eam8.*w allele where the peak phase of *HvCCA1, HvTOC1* and *HvGI* expression was also earlier (Faure *et al*. 2012). Similarly, the same study reported loss of rhythmic oscillations in gene expression upon transition to LL in barley. We found that functional *TtELF3* is required for high amplitude *TtLHY* rhythms, and that *TtELF3* loss of function mutants have increased trough transcript abundance of *TtGI* (Figure 6E). The similarity of these observations to those in barley and rice lines carrying *ELF3* mutations (Zhao *et al*. 2012, Boden *et al*. 2014) suggests that *ELF3* genes might have similar functions across grasses.

Wheat PHYTOCHROME B (PHYB) and PHYC are required for accelerated heading in long days (Chen *et al*. 2014, Pearce *et al*. 2016, Kippes *et al*. 2020, Bouché *et al*. 2022) and PHYB and PHYC are required for the induction of *Ppd-*1 expression during night-breaks in wheat (Pearce *et al*. 2017). In Arabidopsis PHYB co-locates at sites bound by the EC (Ezer *et al*. 2017) and in Brachypodium ELF3 appears to affect PHYC activation of *Ppd-1* physical interaction (Gao *et al*. 2019). Together these results support the hypothesis that wheat ELF3 may act as an integrator of light signalling downstream of the phytochromes and upstream of Ppd-1 (Kippes *et al*. 2020) placing it within one of the wheat photoperiod pathways (Shaw *et al*. 2020) (Figure 8).

### An altered role for the evening complex (EC) in wheat

Binding of the Arabidopsis EC to target promoters requires LUX, as both ELF3 and ELF4 lack DNA binding domains, and it is thought this might explain the similar timing of the transcript accumulation (Nusinow *et al*. 2011). However, in wheat the maximal abundance of *ELF3* and *LUX* transcripts is separated by 12 hours (Figure 6F and G). This difference in the peak time of *ELF3* and *LUX* transcript abundance also occurs in *T. monococcum* (Alvarez *et al*. 2016), rice (Zhao *et al*. 2012) and between *ELF3-like1* and *LUX* in maize, sorghum and foxtail millet (Lai *et al*. 2020). This suggests that temporal separation of *ELF3* and *LUX* expression might be a general feature of cereal circadian oscillators. Wheat also lacks a clear orthologue of *AtELF4* and instead contains *ELF4-like* genes (Calixto *et al*. 2015). *ELF4-like* genes might have a conserved role because expression of a barley *ELF4-like* gene restored circadian rhythms to an Arabidopsis *elf4-2* mutant (Kolmos *et al*. 2009). There is further evidence for conservation of EC function in monocots because Brachypodium ELF3 binds Arabidopsis ELF4 and LUX, and can complement Arabidopsis *elf3-2* mutants (Huang *et al*. 2017). The temporal separation of wheat *ELF3* and *LUX* transcript abundance does not preclude the formation of an EC complex, but could suggest one is not formed, or occurs at a different time in the diel cycle to Arabidopsis. Whether through an EC or not, our data demonstrate that functional *ELF3* and *LUX* are required for robust circadian rhythms in wheat (Figures 6–8).

Whilst it is notoriously difficult to attribute plant promoter *cis*-elements to specific patterns of expression (Harmer and Kay 2005), the difference in timing of *TaELF3* and *TaLUX* expression compared to their Arabidopsis orthologues might arise because of promoter sequence differences. Within 5000 bp upstream of the three *TaELF3* homoeologues transcription start sites the most common motifs are linked to morning phased expression (37 morning, 20 evening and 7 midnight) (Supplemental Table S6). The general composition of the *TaELF3* promoters is similar to *TaLHY* (22 morning, 9 evening and 3 midnight) (Supplemental Table S7) but different to the evening expressed *TaLUX* promoters where the most abundant motifs are associated with evening phased expression (32) compared to morning (23) and midnight (12) (Supplemental Table S8). In particular*, TaELF3* promoters contain Morning Element 1 (ME1) and Morning Element 2 (ME2) motifs, associated with morning phased expression (Michael *et al*. 2008), which are also found in the promoter region of the dawn expressed *TaLHY* homoeologues (Supplemental Table S7) but are absent in the *AtELF3* promoter (Supplemental Table S6). The *TaELF3* promoters also contain further morning-related motifs such as the multiple Hormone up at dawn 1 (HUD1) and Hormone up at dawn 2 (HUD2) elements (Michael *et al*. 2008) which are less frequent in the *AtELF3* promoter (Supplemental Table S6).

### Wheat *ELF3* might be a promising breeding target

*ELF3* might be a suitable target for breeding new lines with heading date better adapted to the local environment in the practice of chronoculture (Steed *et al*. 2021). In other species such as pea, lentil and barley, allelic variation in *ELF3* has permitted the cultivation of these crops at different latitudes (Weller *et al*. 2012, Zakhrabekova *et al*. 2012, Lundqvist 2014). Whilst complete loss of functional *ELF3* in wheat significantly advances flowering time, severely disrupts circadian rhythms and decreases the spikelet number (Alvarez *et al*. 2016), we have demonstrated that altering the function of particular homoeologues of *ELF3* can affect heading without a measurable disruption to the circadian oscillator (Figure 5) and the potential associated yield penalties (Dodd *et al*. 2005), providing some functional copies of *ELF3* are present. It might be possible by combining different alleles of *ELF3* with alleles of other circadian clock genes, to precisely tune flowering time suited to the growing environment. This could be tested by measuring flowering time in selected Cadenza TILLING lines (Krasileva *et al*. 2017) containing mutations in other circadian clock genes as Cadenza contains both the *TaELF3-B1* Ser674Gly SNP and the 1D introgression. In addition to the *ELF3* alleles described in this work, we have also identified further *ELF3* alleles including the absence of *ELF3-A1* in Julius (Supplemental Table S9) highlighting a range of *ELF3* alleles within modern wheat cultivars which could be incorporated into breeding programmes. The variation at *ELF3* might have occurred because of the positioning of *ELF3* within the subtelomeric region (Aguilar and Prieto 2020) which has enhanced levels of recombination which can lead to increased number of sequence and structural variants. This has been observed for other regions, recent studies have reported that a deletion at the distal end of chromosome 4AL is associated with altered heat tolerance (Zhai *et al*. 2021) and a deletion at the distal end of 5AL encompasses the *Rht12* locus which is associated with dwarfing (Sun *et al*. 2019). Secondly, variation at *ELF3* may have been selected by past breeding efforts, leading to a change in allele frequencies, due to advantageous adaptation of flowering time to local climates similar to that seen for selection of *Ppd-1* alleles based on latitudinal cline (Cockram *et al*. 2007) and altitude (Guo *et al*. 2020). A more systematic understanding of circadian clock gene alleles and their combinations will allow for a more targeted approach in optimising flowering time without impacting the functioning of the circadian clock.

## Materials and methods

### Plant material

The eight MAGIC parents; Alchemy, Brompton, Claire, Hereward, Rialto, Robigus, Soissons and Xi-19 were obtained from NIAB (National Institute of Agricultural Botany, UK). The *Ttelf3*-null and *Ttelf3*-WT lines were donated by Jorge Dubcovksy (Alvarez *et al*. 2016) where the *Ttelf3*-null line is the progeny of bulked *elf3-null/Ppd-A1b* and the *Ttelf3*-WT line is progeny of bulked *ELF3WT/Ppd-A1b* (Table 4). By outcrossing we selected lines that contained just the *Ttelf3-A1* mutation (herein *Ttelf3*-Anull) or just the *Ttelf3-B1* mutation (*Ttelf3*-Bnull). The Chogokuwase, Minaminokomugi and Geurumil lines were donated by Hidetaka Nishida (Mizuno *et al*. 2012).

**Table 4:** Wheat lines used in this study. GRU refers to the germplasm resource unit at the John Innes Centre, Norwich.

| Species | Cultivar | Genotype | Reference |
| --- | --- | --- | --- |
| *Ae. speltoides* |  |  | GRU 2140023 |
| *T. aestivum* | Alchemy |  | (Mackay *et al.* 2014) |
| *T. aestivum* | Brompton |  | (Mackay *et al.* 2014) |
| *T. aestivum* | Claire |  | (Mackay *et al.* 2014) |
| *T. aestivum* | Hereward |  | (Mackay *et al.* 2014) |
| *T. aestivum* | Rialto |  | (Mackay *et al.* 2014) |
| *T. aestivum* | Robigus |  | (Mackay *et al.* 2014) |
| *T. aestivum* | Soissons |  | (Mackay *et al.* 2014) |
| *T. aestivum* | Xi-19 |  | (Mackay *et al.* 2014) |
| *T. aestivum* | Cadenza |  | GRU WBCDB0013-PG1 |
| *T. aestivum* | Jagger |  | GRU PANG0006 |
| *T. aestivum* | Spark |  | GRU WBCDB0052 |
| *T. aestivum* | Chinese Spring |  |  |
| *T. aestivum* | Chogokuwase |  | (Mizuno *et al.* 2016) |
| *T. aestivum* | Minaminokomugi |  | (Mizuno *et al.* 2016) |
| *T. aestivum* | Geurumil |  | (Mizuno *et al.* 2016) |
| *T. aestivum* | Axona |  | GRU W3580 |
| *T. aestivum* | Cordiale |  | GRU W10003 |
| *T. aestivum* | Maris Fundin |  | GRU W1357 |
| *T. aestivum* | Tonic |  | GRU W3572 |
| *T. monococcum* |  |  | GRU T1040057 |
| *T. sphaerococcum* |  |  | GRU 1210007 |
| *T. timopheevii* |  |  | GRU 116002 |
| *T. turgidum* | Kronos | *Ttelf3*-null line | (Alvarez *et al.* 2016) |
| *T. turgidum* | Kronos | *TtELF3*-WT | (Alvarez *et al.* 2016) |
| *T. turgidum* | Kronos | *Ttelf3*-Anull | In house |
| *T. turgidum* | Kronos | *Ttelf3*-Bnull | In house |

### Plant growth conditions

Seeds were surface sterilised by washing once in 70% (v/v) ethanol for one minute (Fischer Scientific, UK), once in autoclaved water for one minute, followed by 15 minutes in 10% (v/v) sodium hypochloride solution (Fischer Scientific, UK) and then washed a further three times in autoclaved deionised water. Seeds were sown directly onto 96 well modular trays (P G Horticulture Ltd, UK), containing a 1:1 mix of Levington M2 potting compost (Levington, UK) and vermiculite (fine, Willian Sinclair Horticulture, UK) and treated with the insecticide Intercept 70W (0.02 g L-1 soil, Bayer, Germany). Plants were grown in growth chambers (Conviron, Canada) under long days and constant temperature (16 h L: 8 h D; 200 µmol m^−2^ s^−1^; 20 °C or 16 h L: 8 h D; 400 µmol m^−2^ s^−1^; 22 °C) or in growth cabinets (see specific methods).

### Field trials

In 2012/2013 a fully replicated yield trial of 784 F_5_ MAGIC lines at the NIAB experimental farm in Cambridge, UK was scored by NIAB for the timing of Growth Stages on the Zadok’s scale (Zadoks *et al*. 1974, Mackay *et al*. 2014). Each plot was assessed every two days for its growth stage based on representative plants at the centre of each plot. Once a plot reached GS55 (half of ear emerged above flag leaf ligule) the date was logged. The phenotype was assessed with growth stage measurements scored as days after sowing. In the 2013/2014 field season the same growth stage phenotypes were collected from a fully replicated yield trial of 784 F_6_ MAGIC lines.

### Analysis of field phenotypes

Asreml-R 3.0 (Gilmour *et al*. 1997) was used to minimise or remove spatial effects in phenotype data due to field variation.

Marker genotypes and their respective chromosomal groupings from (Gardner *et al*. 2016) were used. The genetic map was reordered visually using the R package mpMapInteractive (Shah 2013) to improve ordering and consider physical ordering information from IWGSCRefSeqv1. Recombination fractions were recalculated in mpMap (Huang and George 2011) for each chromosome using mpestrf(). Recalculating recombination fractions on a per chromosome basis with the existing marker groupings significantly reduced computation resources required and enables the use of finer recombination bins (r <-c(0:5/600, 2:20/200, 11:50/100) compared to the default values. Genetic map lengths were recalculated using the Kosambi mapping function in computemap().

To reduce computational effort a skimmed genetic map with only unique mapping locations was used to calculate the probability that each location on a genome was inherited from each parent and conduct QTL mapping in MPWGAIM only (Verbyla *et al*. 2014). A marker for each unique coordinate was chosen which had the lowest missing calls and preferably a clear BLAST hit to the IWGSCRefSeqv1 reference sequence, which was used to assign physical mapping coordinates. Out of 18089 mapped markers 4458 were chosen for founder probability calculations.

To test and validate our pipeline and analysis tools we used them to interrogate a data set describing flowering time in a 2011/2012 nursery of the NIAB eight Parent MAGIC population that had previously been analysed using Bayesian Networks (Scutari *et al*. 2014). Reanalysis of this data using MPWGAIM (Verbyla *et al*. 2014) detected 13 significant QTLs for flowering time with significant founder effects (Supplemental Table S1).

This analysis identified a major QTL (18.3% genetic variation explained, LOGP 18.33) between the markers Kukri_c27309_590 (48.57 cM) and BS00064538_51 (56.64 cM), defining a maximum mapping interval of 31.81 Mb – 34.23 Mb between the two mapping bins as several markers map to the 56.64 cM bin. The mean founder effects of the QTL predict a significant acceleration of flowering time for Soissons, which is the only parent carrying the photoperiod insensitive *Ppd-D1a* allele (Bentley *et al*. 2013) and furthermore *Ppd-D1* maps within the QTL (33.95 Mb). This analysis of the effect of *Ppd-D1a* on flowering time validated MPWGAIM and our QTL mapping pipeline in MAGIC.

### Synteny analysis between Chinese Spring and Jagger 1D

A conserved haplotype block between Cadenza and Jagger at the distal end of chromosome 1D was identified using the Crop Haplotypes web browser (Crop Haplotypes (crop-haplotypes.com) (Brinton *et al*. 2020). Physical locations of genes on the Chinese Spring 1D chromosome (IWGSCRefSeqv1.1) and Jagger 1D chromosome (PGSBv2.0) were mapped using the mapviewer v2.6.1 (https://10wheatgenomes.plantinformatics.io/mapview) and then confirmed manually using EnsemblPlants.

### SNP mining and analysis

820k and 35k SNP array data was obtained from the CerealsDB website (Wilkinson *et al*. 2012) and the 1000Exome data from the 1000 Exomes project website (http://wheatgenomics.plantpath.ksu.edu/1000EC/). Mapping of 35k and 820k SNPs to the IWGSCv1.1 Ref sequence were obtained from EnsemblPlantsv51. SNPs for the Cadenza/Jagger haplotype (chr1D: 482.3Mb – end of chromosome 1D) were extracted and heterozygous calls set as missing. Distance matrices and phylogenetic trees were constructed using the R package “ape” (Paradis and Schliep 2019) 100 bootstrap permutations were run also using the R package “ape”.

To investigate the prevalence of the Ser674Gly SNP in a broad range of germplasm we analysed variation data from multiple sources including the 820k SNP genotyping array (Winfield *et al*. 2016), the wheat HapMap panel (Jordan *et al*. 2015), the 1000 wheat Exome project (He *et al*. 2019) and the 10+ Wheat Genome project (Walkowiak *et al*. 2020). We found that the Ser674Gly SNP was also present in the HapMap panel as chr1B_685645813 and within the 1000 Exome data as scaffold48561_337271.

### Phylogenetic tree construction

*ELF3-D1* orthologous groups were retrieved from public repositories. Coding sequences were aligned using the MCoffee aligner on default settings (https://pubmed.ncbi.nlm.nih.gov/16556910/). Maximum Likelihood (ML) Phylogenetic trees were build using CLC Genomics Workbench 21.0.3. The best substitution model was determined using the Model testing function before running Molecular Phylogenetic analysis by Maximum Likelihood method with 100 permutations for Bootstrap values. The *ELF3* gene from *Brachypodium distachyon* was used as an outgroup.

### Quantitative Real-Time PCR

The *TtELF3*-WT and *Ttelf3*-null lines were sown and grown in two antiphased LED growth chambers (Conviron) under long day conditions (16h L: 8h D; 250 µmol m^−2^ s^−1^, 20°C day: 16°C night). Sampling commenced 16 days post-sowing, the first true leaf was sampled every 3 hours for 96 hours. After the first 24 hours of sampling the cabinet was switched to constant light and temperature (250 µmol m^−2^ s^−1^, 20°C). Total RNA was extracted from leaf samples using the KingFisher Flex purification system (Thermofisher scientific) in conjunction with the Maxwell® HT simplyRNA Kit (Promega). RNA concentration was determined using the Nanodrop ND-1000. DNA contamination was removed using the Arcticzyme HL-DNAse kit. RNA integrity was checked using the Fragment AnalyzerTM (AATI, USA) before cDNA was synthesised from extracted RNA using the High-Capacity cDNA reverse transcription kit (Applied Biosystems). Three technical replicates of gene specific products were amplified in 10 µl reactions using Power SYBR Green Master mix on a CFX384 Touch Real-Time PCR detection system. Gene expression levels were determined relative to the expression of two housekeeping genes, *RP15* and *RPT5A,* that were selected using the GeNorm alogorithm (Vandesompele *et al*. 2002) (Table 5).

**Table 5:**
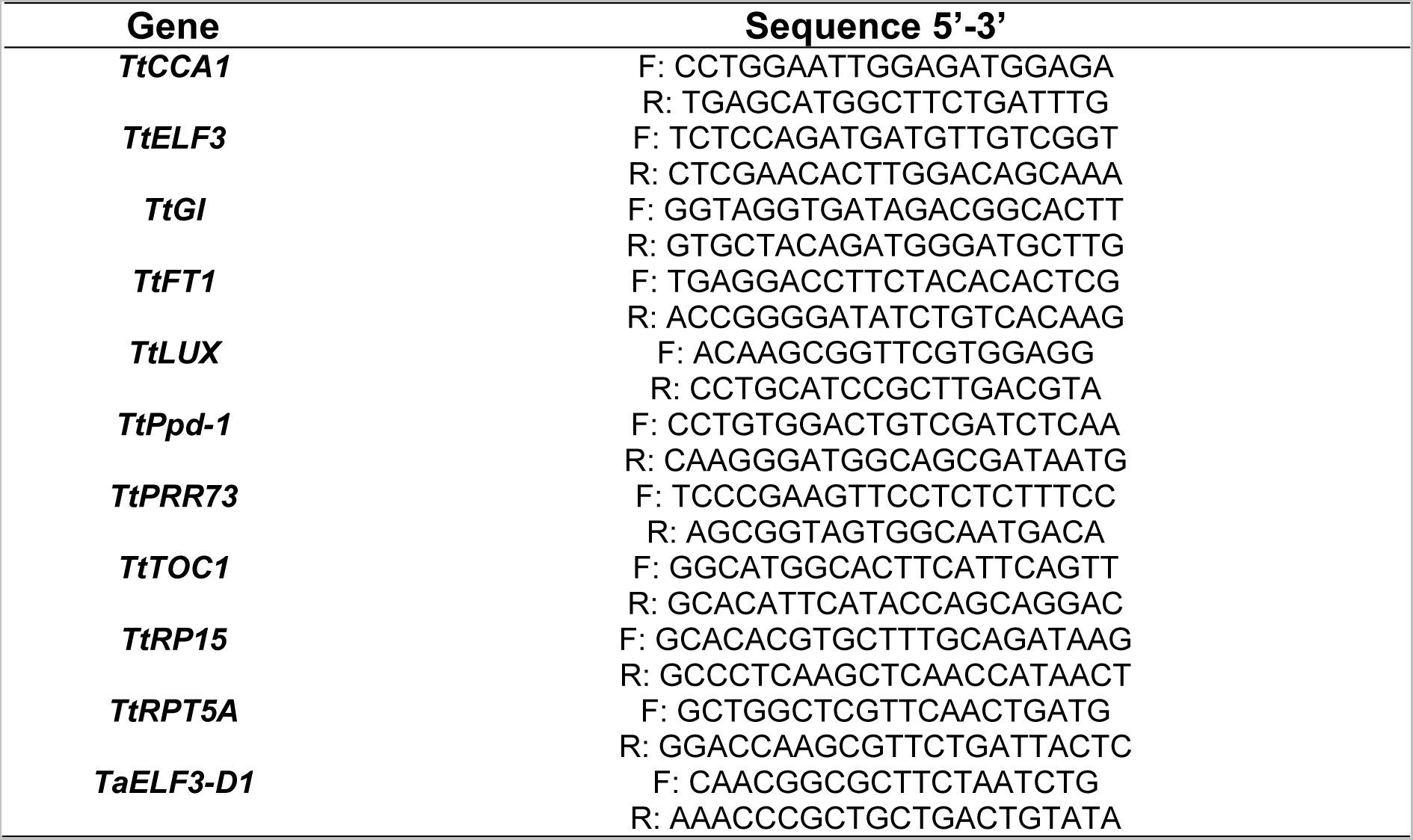
Primers used in this study

For quantification of *TaELF3* transcript abundance Cadenza, Chinese Spring, Claire, Paragon and Robigus seed was sown and grown in a Conviron growth room (16h L: 8h D; 400 µmol m^−2^ s^−1^, continuous 22°C. After 14-days leaf samples were taken and RNA extracted using the RNAeasy plant mini kit (Qiagen, UK) with an on column DNAse digest (Qiagen, UK). RNA concentration was determined on the Nanodrop ND-1000 and cDNA synthesised from 500 ng RNA using the RevertAid First Strand cDNA synthesis kit (Thermo Scientific, UK). Technical duplicates of gene-specific products were synthesised in 10 µl reactions using the Rotor-Gene SYBR Green PCR kit (Qiagen, UK) on a Rotor-Gene 6000 Real-Time PCR machine (Qiagen, UK).

### PCR of *ELF3-D1*

DNA extracted using the DNeasy plant mini extraction kit (Qiagen, UK). The presence of *ELF3-D1* was confirmed by PCR using the *TaELF3-D1* primers listed in Table 5. PCR was performed using the DreamTaq Green PCR kit (Thermo Scientific, UK) on a ProFlex PCR system (Thermo Scientific, UK). PCR reaction conditions were as follows: 3 minutes at 95°C; 40 cycles of 30 s at 95°C, 30 s at 68°C, 90 s at 72°C; 15 minutes at 72°C.

### Delayed chlorophyll fluorescence and prompt chlorophyll *a* **fluorescence imaging**

Leaf samples were taken from the upper half of the first true leaf of 16-day old seedlings (non-vernalised) grown and entrained in Conviron growth cabinets and were cut into 3 mm by 3 mm pieces and placed on solid 0.8% (w/v) bactoagar, 0.5 x MS Vitamin medium (M3900, Sigma-Aldrich, UK) supplemented with 0.5 µM benzyl-aminopurine (BAP), poured in a black 96 well imaging plates (Greiner, UK). Delayed fluorescence (Gould *et al*. 2009) was measured for 60 s in an LB985 Nightshade (Berthold, UK) camera. Between measurements of DF light was supplied from blue (470 nm) red (660 nm) and LED arrays at 100 µmol m^−2^ s^−1^ during the experiment. Temperature was maintained at 20°C.

Chlorophyll *a* fluorescence and calculations of parameters were carried out using a CFimager (Technologica Ltd, Colchester, UK) and the image processing scripts provided by the manufacturer. Chlorophyll fluorescence images were captured using a Stingray F145B ASG camera (Allied Vision Technologies, UK) through an RG665 long pass filter to exclude blue light from the LEDs. When imaging under continuous light, leaf fragments were exposed to 40 minutes of 100 µmol m^−2^ s^−1^ of blue light followed by a saturating pulse of 6172 µmol m^−2^ s^−1^ blue light for 800 ms then 20 minutes of darkness (with non-actinic measuring light on) followed by a second saturating pulse of 6172 µmol m^−2^ s^−1^ for 800 ms.

Circadian measures of amplitude, phase, period and relative amplitude error were calculated using the FFT-NLLS method in Biodare2 (www.biodare2.ed.ac.uk) (Zielinski *et al*. 2014). Where necessary data was normalised to between −1 and 1 using the normalisation feature of Biodare2.

## Heading date measurements in *TtELF3* kronos lines

Seeds sown as described above and moved to long day conditions (Conviron growth cabinet, 16 h light: 8 h dark; 400 µmol m^−2^ s^−1^, 22°C). After two weeks seedlings transplanted to 10 cm pots. Heading (GS55) was scored once per day.

### Direct contact leaf temperature measurements

Leaf surface temperature relative to background temperature was measured using an in-house built device and provides an alternative measure of circadian rhythms caused by circadian regulated changes in the stomatal aperture (Dakhiya and Green 2019). The temperature logger was validated by measuring circadian rhythms of “in-phase” and “anti-phase” wheat leaves where peak temperature, and therefore maximum closure, occurred during the subjective night (Supplemental Figure S15). Leaf temperature measurements were taken from fully expanded flag leaf of 4-week-old plants under continuous light and temperature (200 µmol m^−2^ s^−1^, 20°C) that were previously grown under long days (16h L: 8h D; 200 µmol m^−2^ s^−1^, 20°C day: 15°C night) in a growth cabinet (Conviron, UK).

Wheat was grown in 10 cm pots. Circadian measurements of leaf temperature were taken when the pants were four-weeks old. Type-K thermocouples were attached to the leaf such the end of the thermocouple was pressing onto the surface of the fully expanded flag leaf. This was kept in position by positioning a paperclip onto the leaf and fixing the paperclip to a stake. The thermocouples were connected to an Arduino Mega using the Adafruit 1778, a 10-bit D-A converter and x20 amplifier giving a resolution of ± 0.1°C with a DS3231 RTC added to keep time, temperature was logged every 30 s. Background temperature was measured using unattached thermocouples. Leaf temperatures were averaged using a 60-point moving average, background temperature was then subtracted from leaf temperature to give leaf temperature relative to background.

## Acknowledgements and Funding

LW, GS and LT were supported by BBSRC studentships BB/K011790/1, BB/M015416/1 and BB/M011194/1 awarded to MAH and AARW, GS is currently supported by UKRI BBSRC grant BB/S006370/1 awarded to AARW. DCR is supported by UKRI BBSRC grant BB/S002251/1 and GP-C is supported by UKRI BBSRC grant BB/W001209/1, both awarded to AARW.

## Contributions

AARW, MAH, AG, KG, LW and GS conceived this study and planned the experiments. LW, GS and LJT performed the experiments and analysed data. DCR and GP-C assisted with scoring heading date and sampling for qRT-PCR. AARW, LW and GS wrote the first draft of the manuscript. AARW, MAH, KG, LW, GS and LJT contributed to the revision of the manuscript.

