## Supplemental Figures and Tables for "Wheat *EARLY FLOWERING3* is a dawn-expressed circadian oscillator component that regulates heading date"

**Supplemental Material**

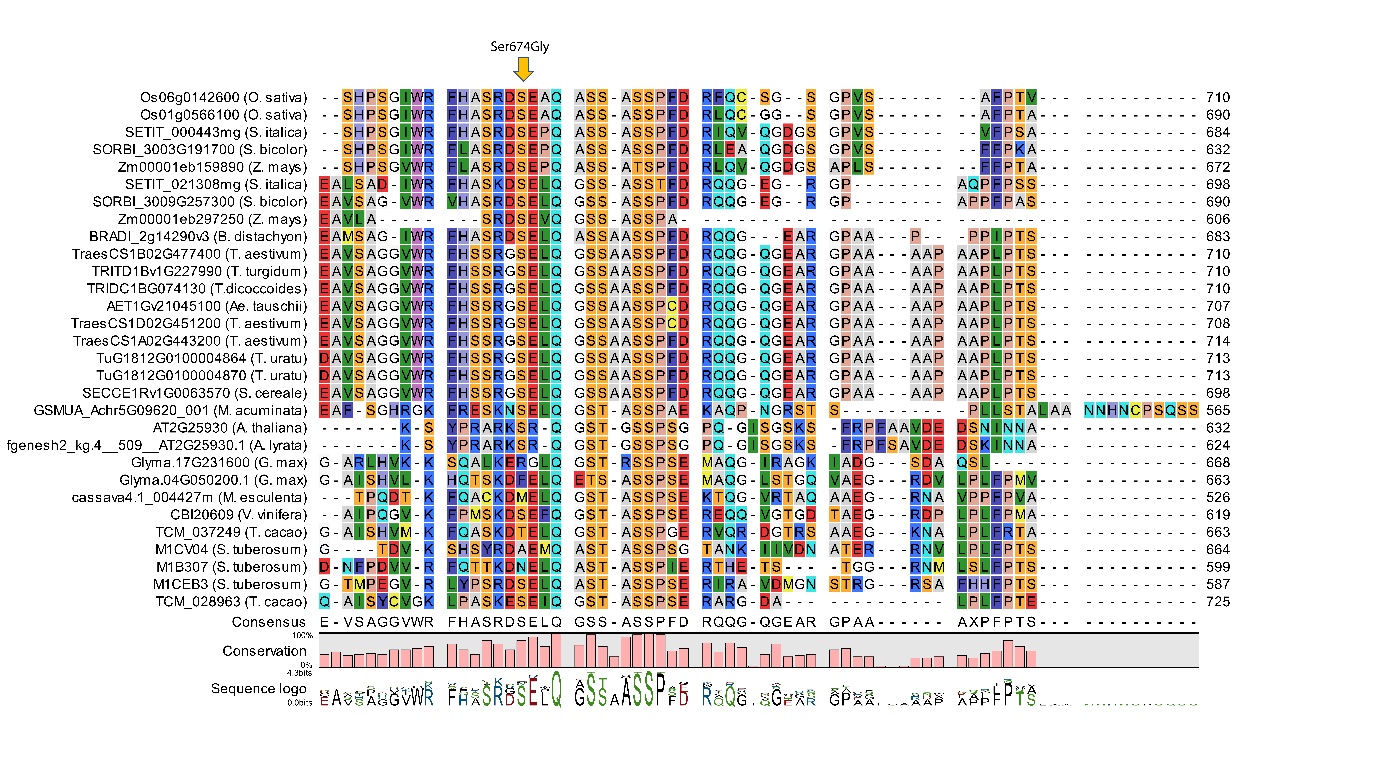

**Supplemental Figure S1: ELF3 Ser674 is highly conserved in monocots.** MCoffee alignment visualised in CLC Genomics Workbench, Amino acids coloured based on RasMol colour scheme.

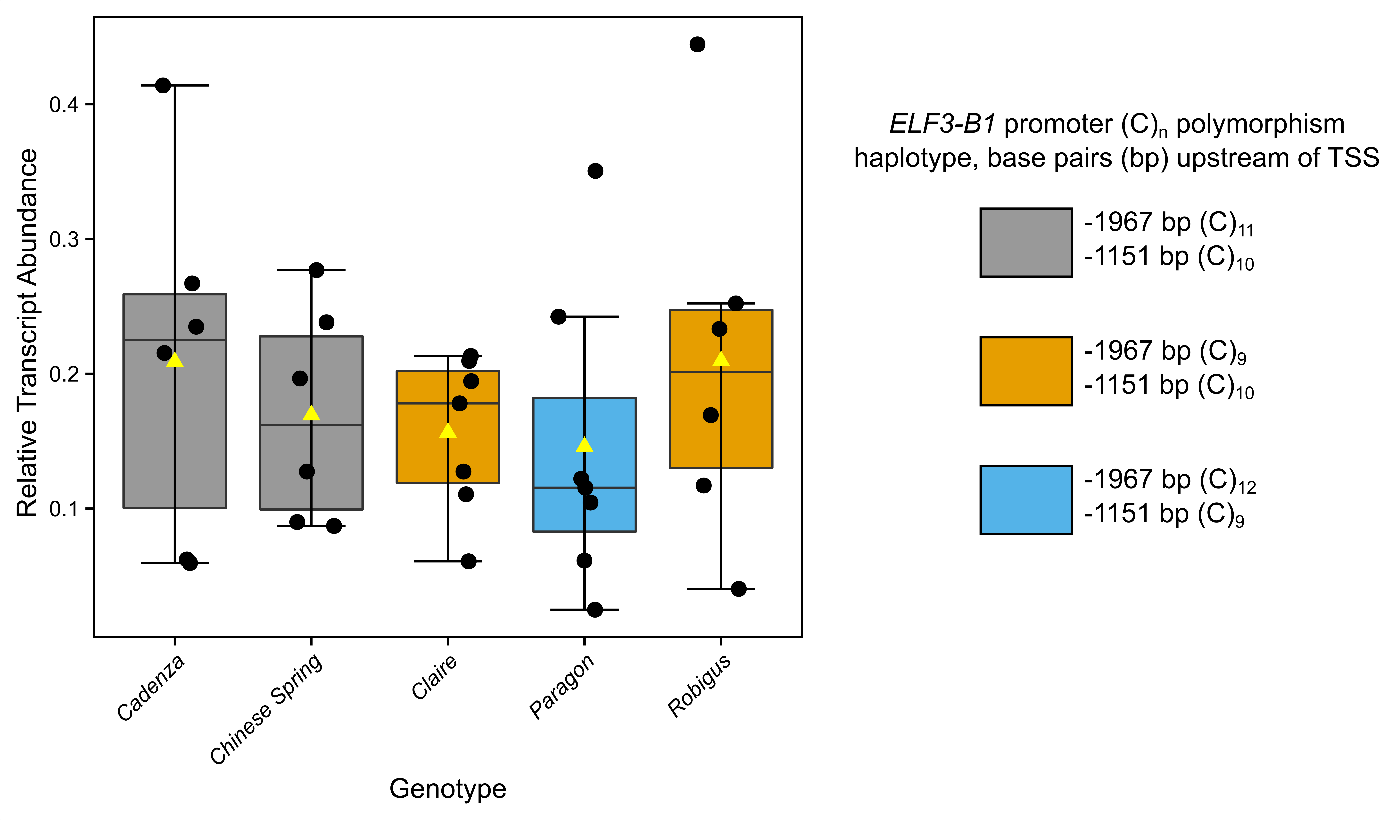

**Supplemental Figure S2: Relative expression of *TaELF3* in five wheat cultivars that differ in *TaELF3-B1* promoter (C)_n_ polymorphic *loci.*** Expression of *TaELF3* relative to *RP15* and *RPT5A* at dawn. Upper and lower hinges represent the first and third quartiles (25^th^ and 75^th^ percentiles), the middle hinge represents the median value, yellow triangle represents mean value and black dots represent individual replicates.

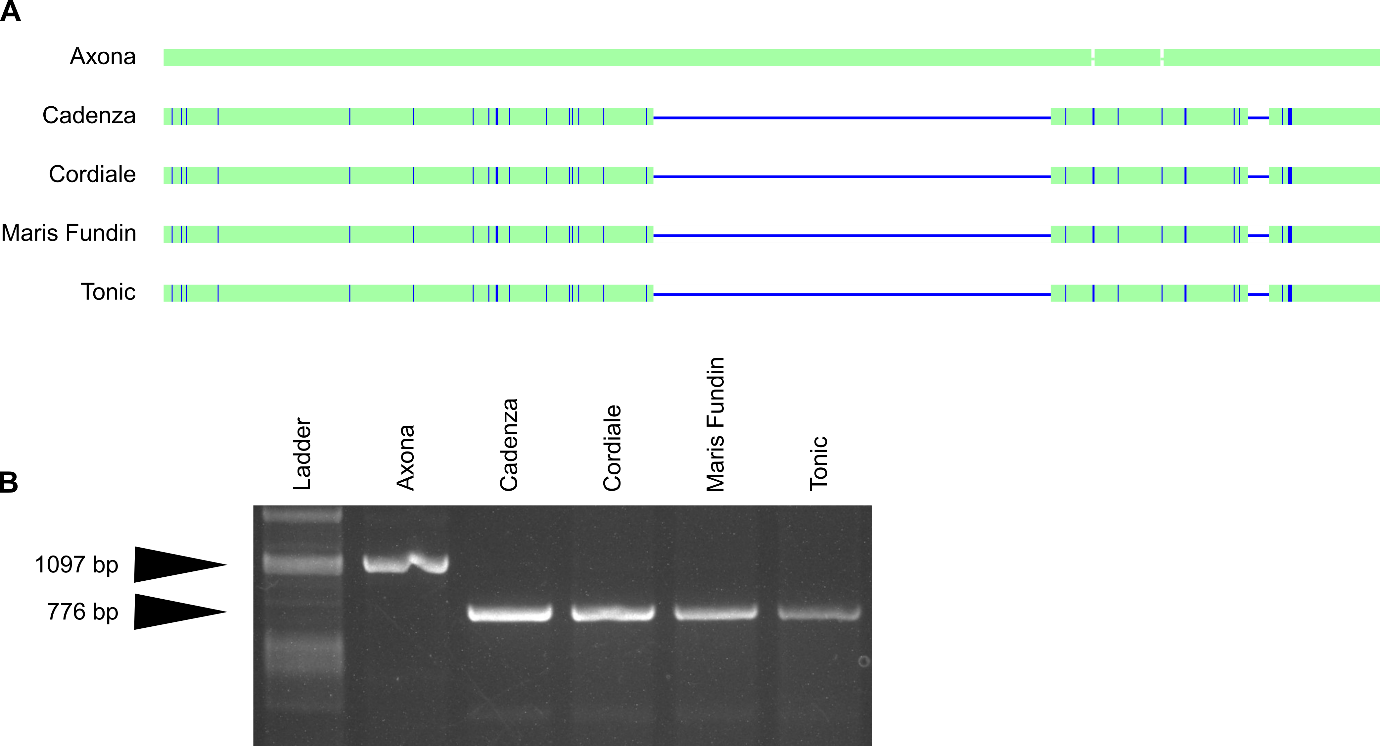

**Supplemental Figure S3: The Cadenza/Jagger *ELF3-D1* intronic deletion is present within Cordiale, Maris Fundin and Tonic*.***

(A) Sequence alignment for PCR fragments between exon 2 and 3 of *ELF3-D1* (see Fig 3) and (B) Agarose gel separation of PCR products. A PCR product at 776bp is indicative of the presence of the introgressed and inverted region.

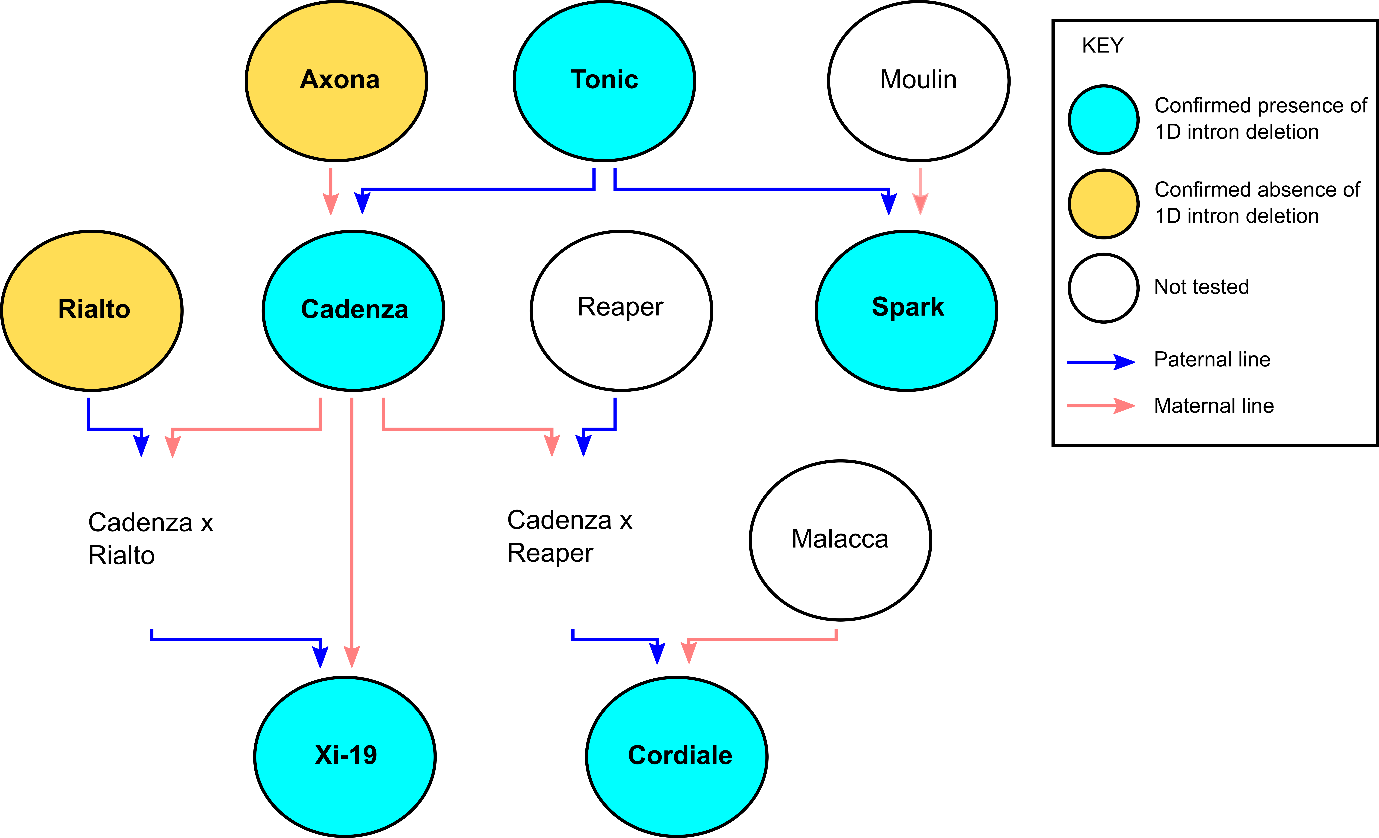

**Supplemental Figure S4: Simplified pedigree diagram shows the relationship between different wheat varieties that carry the Jagger/Cadenza *TaELF3-D1* allele.**

Pedigree diagram constructed from relationships described at <https://wheatpedigree.net> and (Fradgley et al., 2019). Note Cadenza x Rialto and Cadenza x Reaper are not available as germplasm for testing.

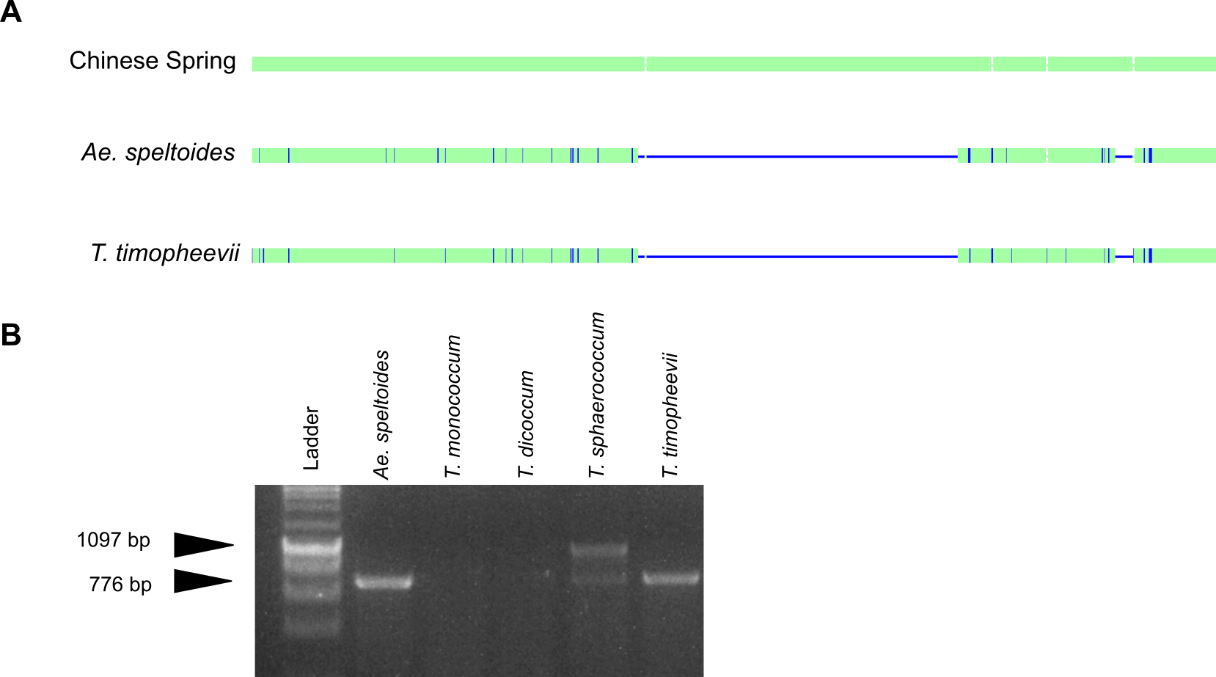

**Supplemental Figure S5: The Cadenza/Jagger *ELF3-D1* intronic deletion is present within *Ae. speltoides* and *T. timopheevii.***

(A) Sequence alignment for PCR fragments between exon 2 and 3 of *ELF3-D1* (see Fig 3) and (B) Agarose gel separation of PCR products. A PCR product at 776bp is indicative of the presence of the introgressed and inverted region.

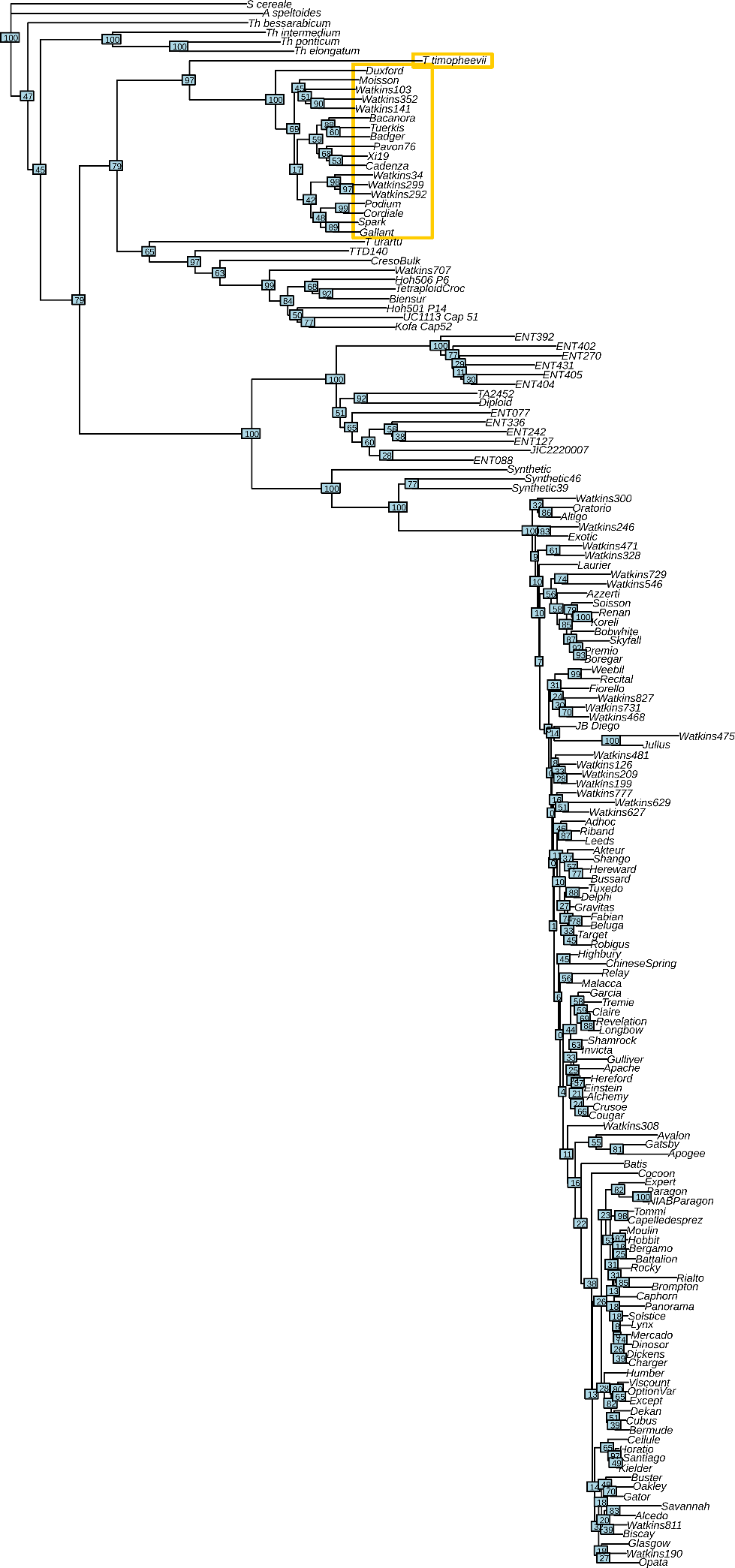

**Supplemental Figure S6: Phylogeny of the 1D Jagger/ Cadenza shared haplotype using the 820k SNP dataset from CerealsDB.**

Neighbour Joining Tree and bootstraps created from 820k SNPs using R package “ape”. All SNPs mapping to the CS coordinates (1D:482.83Mb until 1D end) of the shared Cadenza and Jagger haplotype were used. Heterozygotes were set as missing. Yellow boxes highlight varieties associated with the 1D introgression.

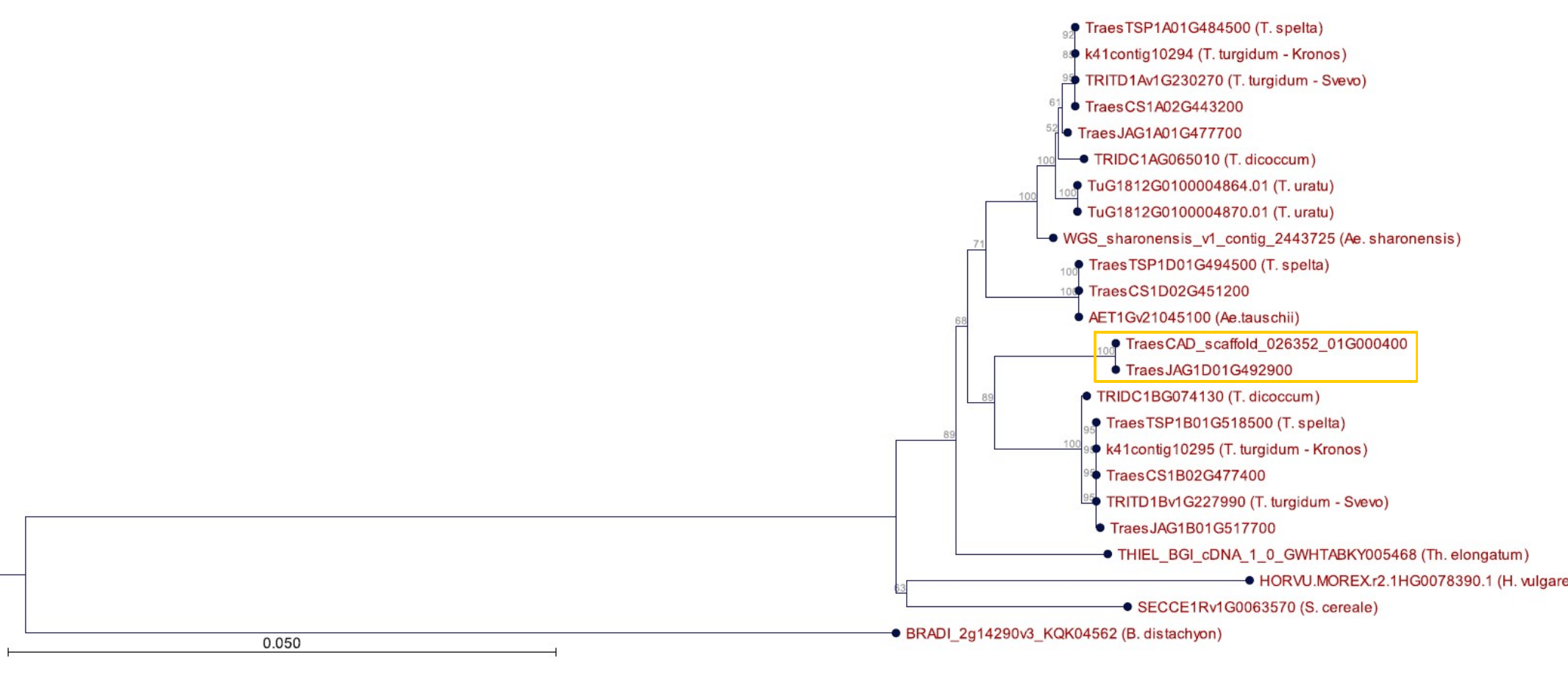

**Supplemental Figure S7:** **Phylogeny of *ELF3-D1* gene.**

Maximum Likelihood phylogeny based on T-coffee alignment of publicly available coding sequences. Cadenza and Jagger highlighted by yellow box.

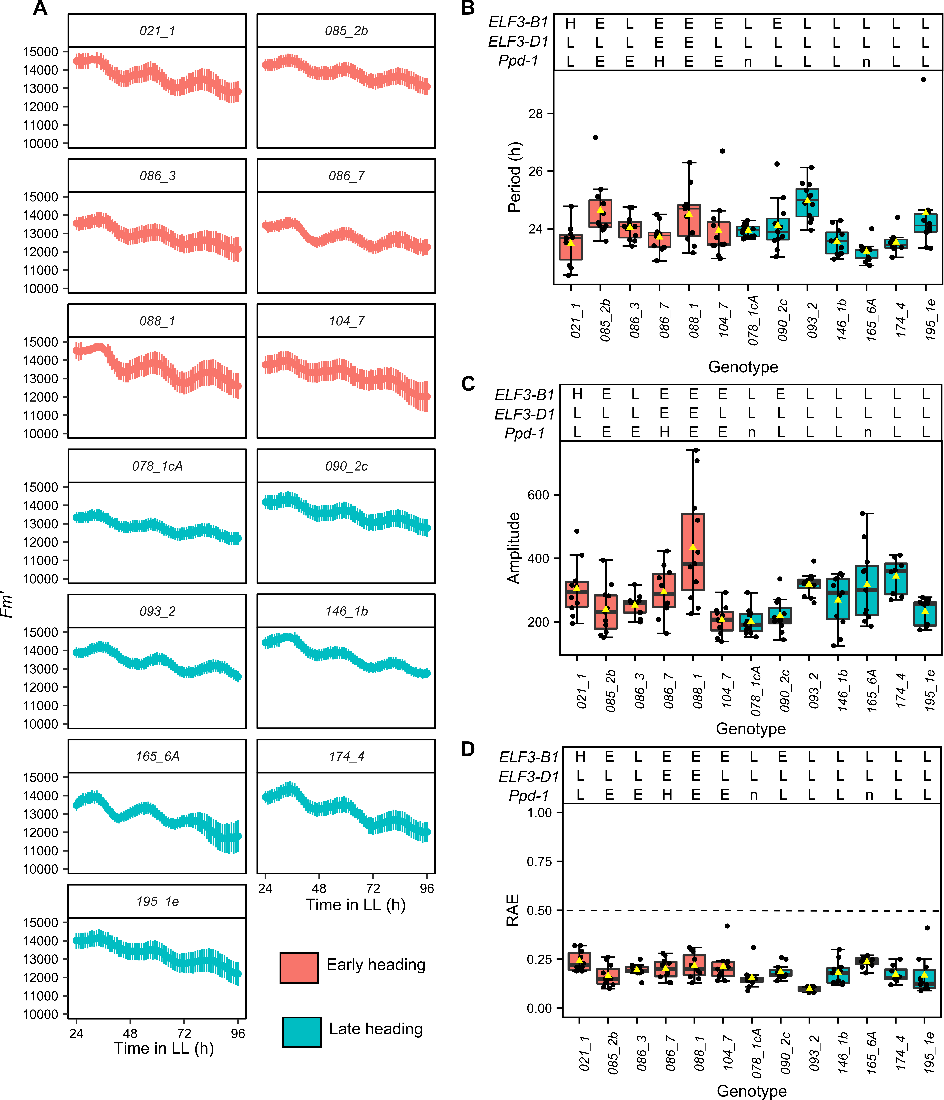

**Supplemental Figure S8: Circadian rhythms of chlorophyll *a* fluorescence in early and late heading MAGIC RILs are not associated with genotype.**

(A) Mean *Fm’*  in continuous light for a selection of early heading (red) and ate heading (cyan) MAGIC RILs from 2014 with error bars representing SEM (n = 12). (B) circadian period (C) circadian amplitude and (D) RAE. (B-C) calculated using FFT-NLLS (Biodare2), upper and lower hinges represent the first and third quartiles (25^th^ and 75^th^ percentiles), the middle hinge represents the median value, yellow triangle represents mean value and black dots represent individual replicates. Top of (B-D) represents the alleles of *ELF3-B1, ELF3-*D1 and *Ppd-*1 where E = early heading associated allele, L = late heading associated allele, H = heterozygous and n = no sequencing information. Horizontal dashed line in (E) represents rhythmic cut-off where RAE < 0.5 is considered rhythmic.

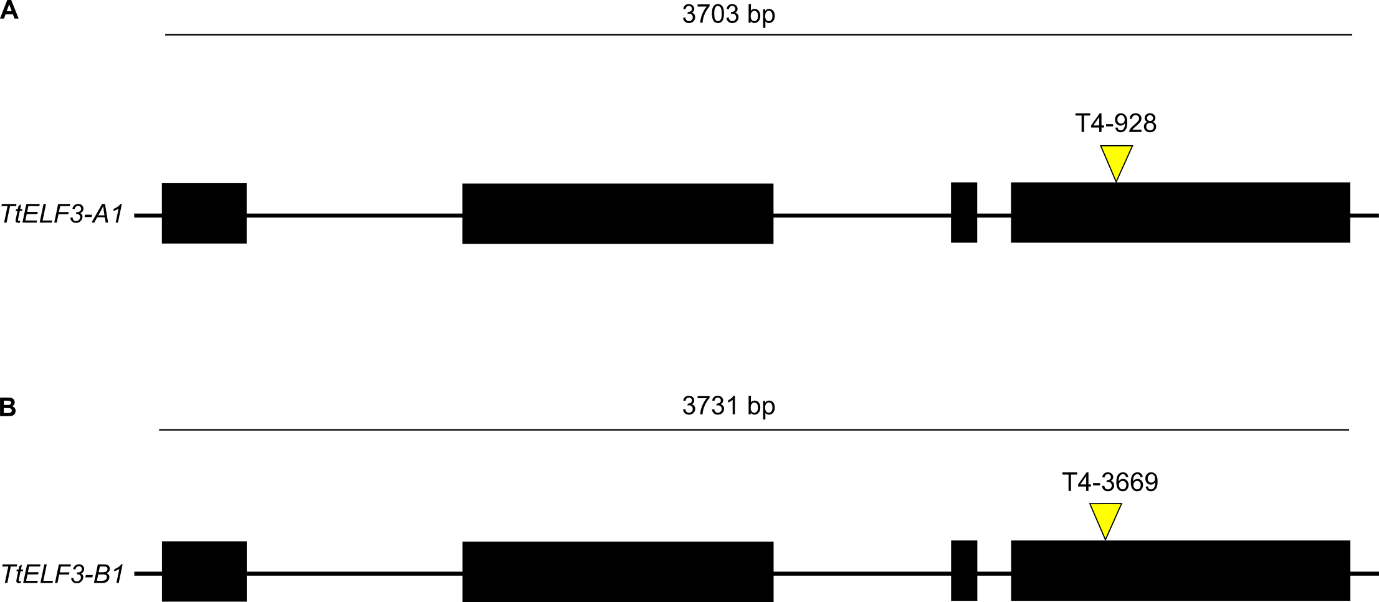

**Supplemental Figure S9: The location of mutations within the A and B sub-genome copies of *TtELF3***.

(A-B) Scale diagram of (A) *TtELF3-A1* and (B) *TtELF3-B1* where black rectangles represent exons ordered 5’-3’ (left-to-right), horizontal line above represents length of gene from start to stop codon (bp), space between exons represents introns. Yellow triangles denote the position of premature stop codons denoting the source TILLING line as annotated in Alvarez *et al.* 2016.

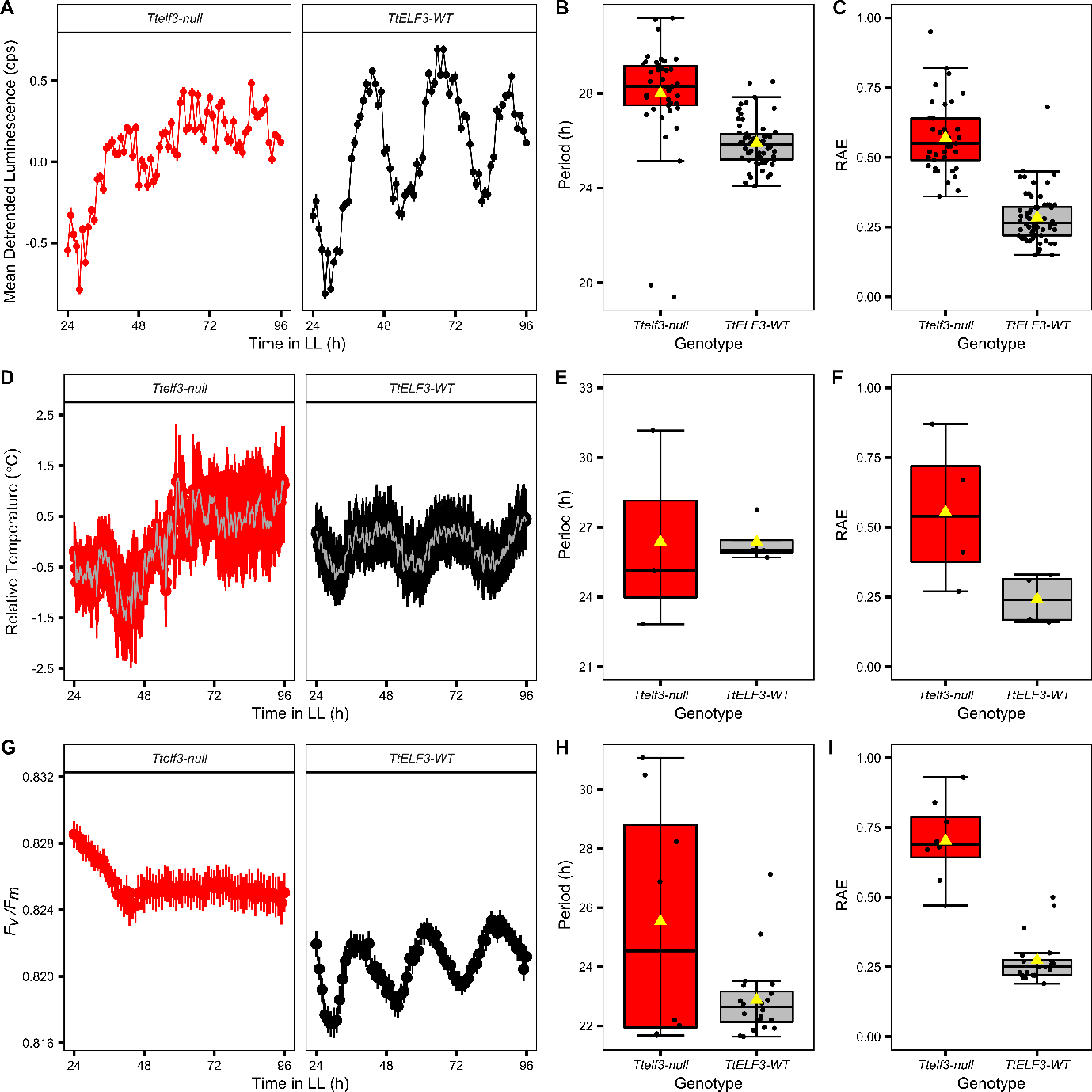

**Supplemental Figure S10: *Ttelf3-null* is arrhythmic for DF and relative leaf temperature in LL which is not a result of leaf stress.**

(A) Mean DF luminescence in counts per second (cps) normalised to -1 to 1 using Biodare2 with error bars representing SEM (n = *TtELF3-*WT 63, *Ttelf3-*null *72*). (B) Amplitude and (C) RAE for *TtELF3-*WT (black) and *Ttelf3-*null (red) in (A) calculated using FFT-NLLS fit in Biodare2 where upper and lower hinges represent the first and third quartiles (25^th^ and 75^th^ percentiles), the middle hinge represents the median value, yellow triangle represents mean value and black dots represent individual replicates. (D) mean leaf surface temperature relative to background and normalised to the mean in LL and constant environmental temperature (20°C) *TtELF3-*WT (black) and *Ttelf3*-null (red) with SEM bars (n = 7), for clarity every 20^th^ point plotted. (E) Circadian period and (F) RAE for plot (D), presented and calculated as for (B-C). (G) Mean *F_v_/F_m_* of *TtELF3-*WT (black) and *Ttelf3*-null (red) in LL (n = 24.) (H) Circadian period (hours) and (C) RAE values for plot (G), calculated as for (B-C).

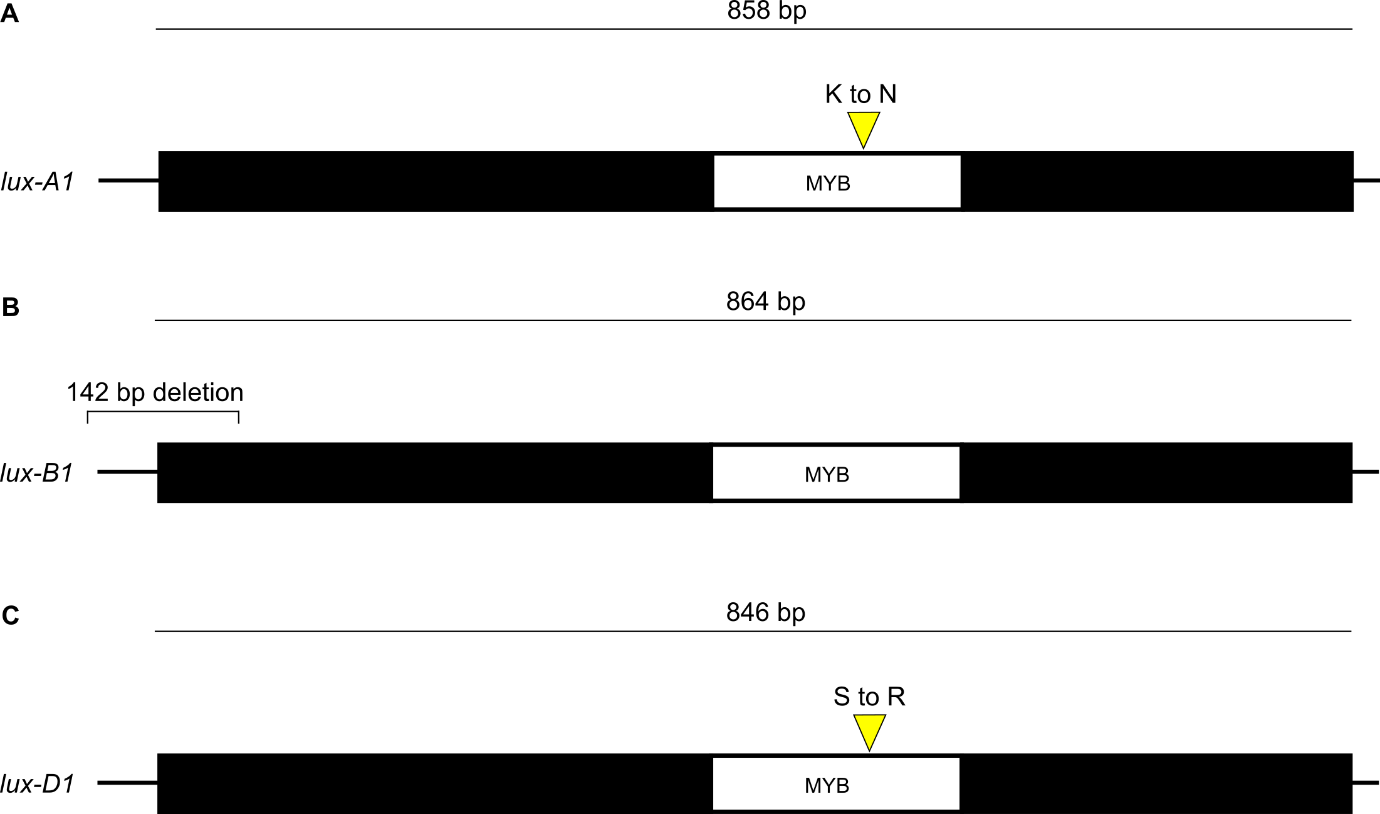

**Supplemental Figure S11: The location of mutations in the different sub-genome copies of *LUX.***

(A-C) Diagram of the A sub-genome (A), B sub-genome (B) and D sub-genome (C) copies of *LUX* where black rectangle represents the *LUX* exon drawn 5’ to 3’ (left-to-right) and the embedded white rectangle denotes the highly conserved MYB-domain. Yellow triangles denote the location of amino acid substitution within the SHAQKYF motif (A, C). (B) line and annotation show the location of 142 bp deletion in the *lux-B1* allele (Mizuno *et al*. 2016)

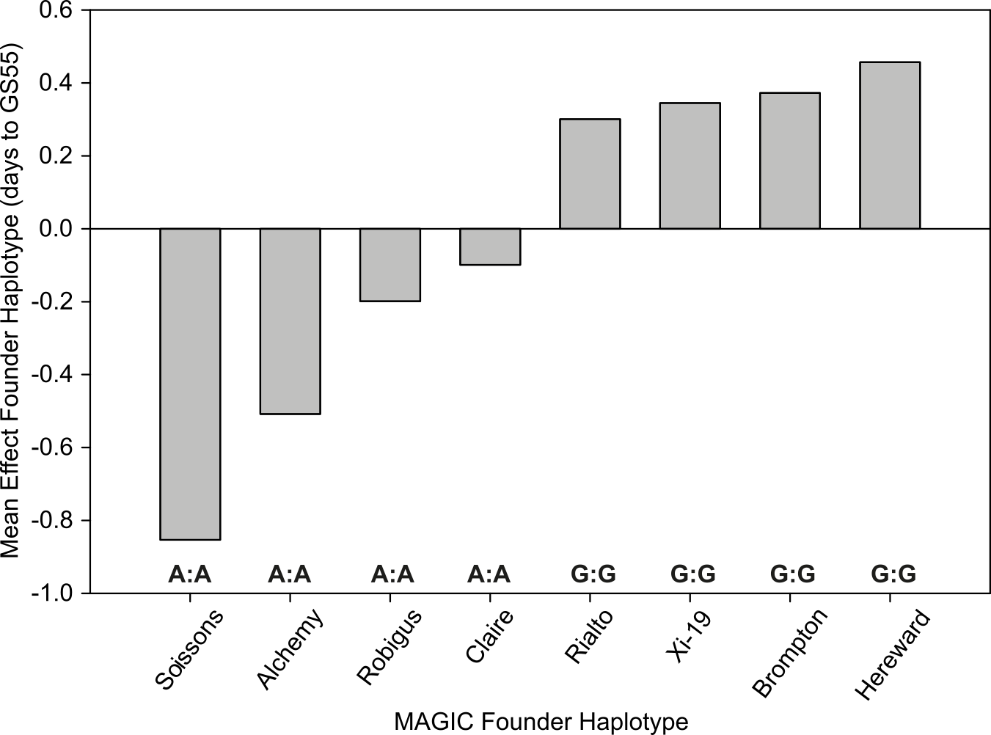

**Supplemental Figure S12: The Ser674Gly SNP is a candidate for the causal *Eps-B1* SNP**.

KASP Genotype association with MAGIC founder haplotype effects at the *Eps-B1* locus. Founder haplotype effects are calculated as the mean of the 2013 and 2014 founder effect. Genotype of A|A corresponds to homozygous for adenine, encoding the ancestral Serine, whilst G|G corresponds to homozygous guanine encoding the non-ancestral Glycine residue.

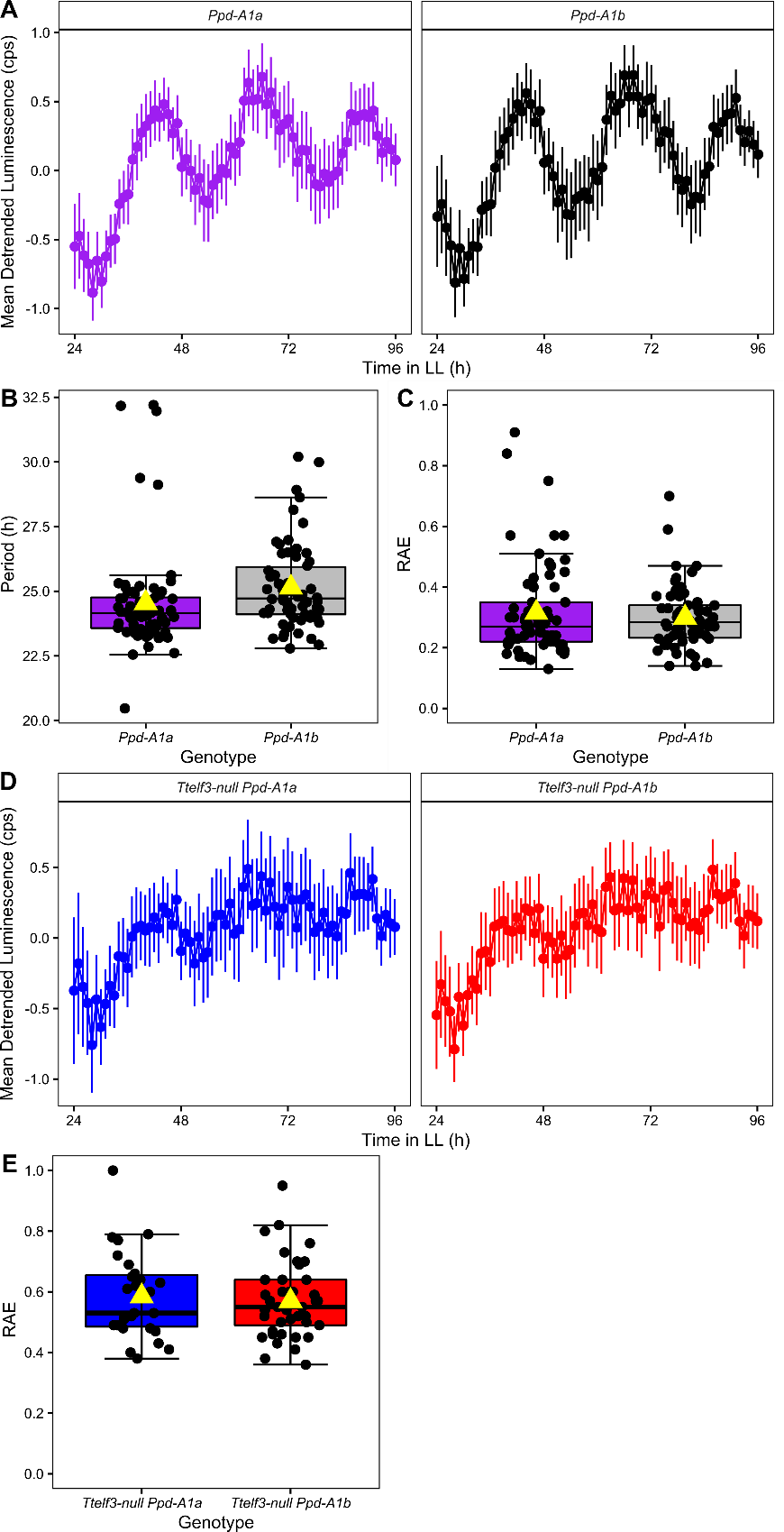

**Supplemental Figure S13: There is no significant difference between the circadian rhythms of kronos and *Ttelf3-null* lines containing the *Ppd-A1a* and *Ppd-A1b* alleles.**

(A) Mean DF luminescence in counts per second (cps) normalised to -1 to 1 using Biodare2 with error bars representing standard deviation (n = *Ppd-A1a* 68, *Ppd-A1b* 62). (B) Amplitude and (C) RAE for *Ppd-A1a* (purple) and *Ppd-A1b* (black) in (A). (D) Mean DF luminescence in counts per second (cps) normalised to -1 to 1 using Biodare2 with error bars representing standard deviation (n = *Ttelf3-null* *Ppd-A1a* 66, *Ttelf3-null Ppd-A1b 2*2) and (E) RAE for *Ttelf3-null* *Ppd-A1a* (blue) and *Ttelf3-null* *Ppd-A1b* (red). Amplitude and RAE calculated using FFT-NLLS fit in Biodare2 where upper and lower hinges represent the first and third quartiles (25^th^ and 75^th^ percentiles), the middle hinge represents the median value, yellow triangle represents mean value and black dots represent individual replicates for plants where estimation was possible.

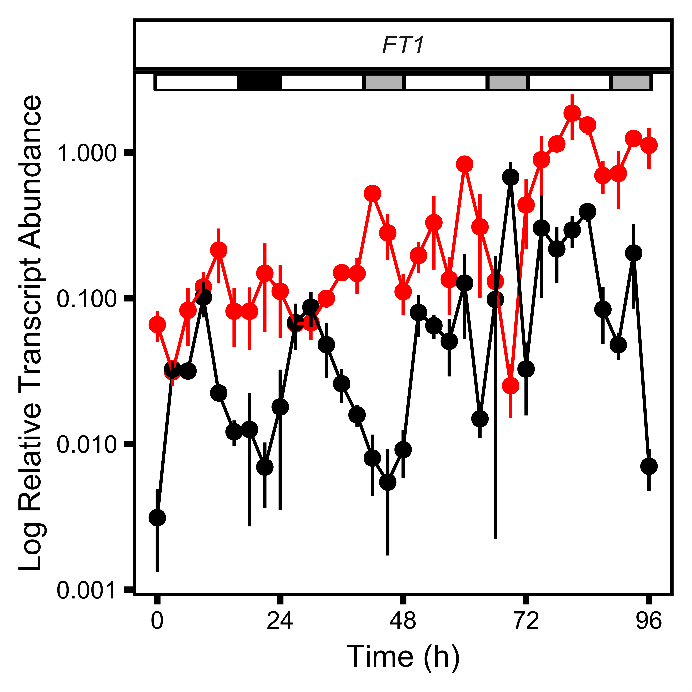

**Supplemental Figure S14: Relative *FT1* transcript abundance in *Ttelf3-*null and *TtELF3-*WT in LD and LL**

Mean *FT1* transcript abundance with SEM bars (n = 3-5) relative to *TtRP15* and *TtRPT5A* expression in *TtELF3-*WT (black) and *Ttelf3-*null (red) in a 24 h LD cycle in long day conditions (16 h light at 250 µmol m^-2^ s^-1^, 20°C: 8 h dark 16°C) followed by constant light and temperature (20°C) from time 24 to 96 h. Horizontal white rectangle represents light, black rectangle represents darkness and grey rectangle represents subjective night.

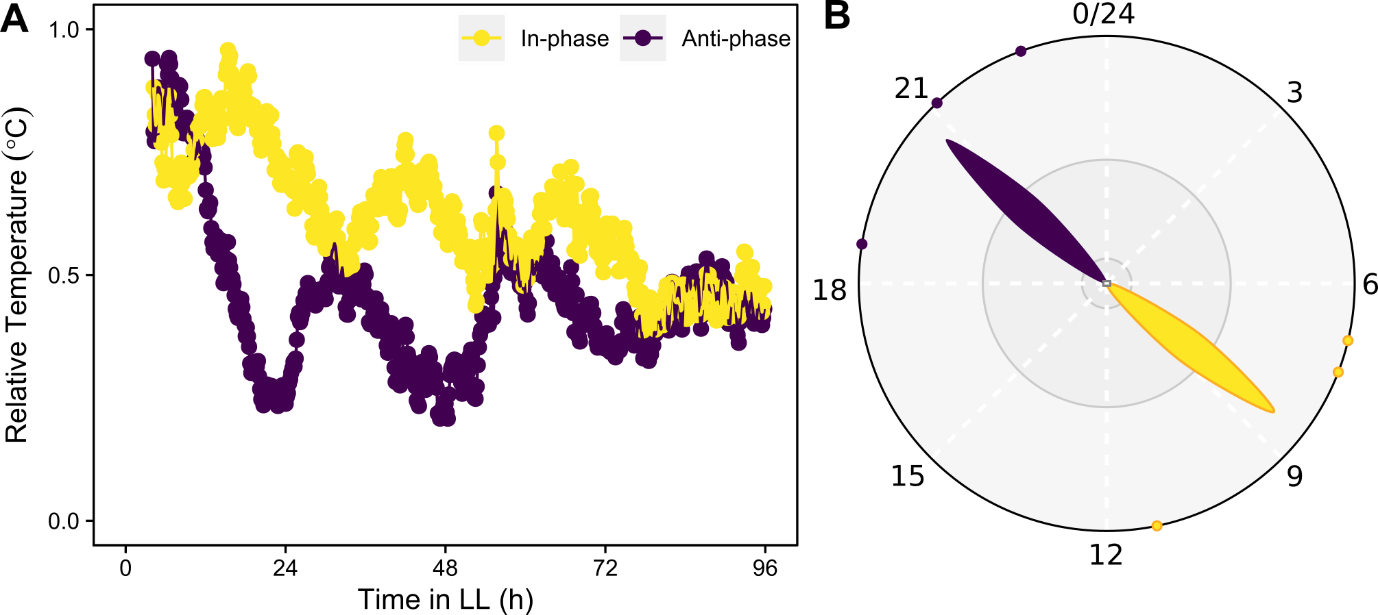

**Supplemental Figure S15: In-phase and anti-phase leaf temperature rhythms in continuous light.**

(A) Mean leaf temperature relative to background (n = 3) of four week old cadenza plants entrained to 16 h light:8 h dark cycles in-phase (yellow) or 16 h light: 8 h dark cycles that were 12 hours anti-phased (purple). (B) Phase plot of data shown in (A).

**Supplemental Table S1: GS55 2012 QTLs**

Output from the reanalysis of the NIAB 2012 GS55 data using MPWGAIM, originally analysed using Bayesian Networks (Scutari et al., 2014). Dist (cM) refers to the markers genetic mapping distance in centiMorgan. Founder corresponds to founder variety in alphabetical order (1 Alchemy, 2 Brompton, 3 Claire, 4 Hereward, 5 Rialto, 6 Robigus, 7 Soissons and 8 Xi-19). Size: Size of effect of each founder allele (days to GS55). LOGP is –log10(p) and corresponds to the overall significance of the QTL as a measure of its strength. Prob: probability QTL is from all founders shown (p). Founder LOGP and Founder Prob are LOGP and Prob for individual founders. % var is the percentage of genetic variance explained by each QTL.

| Chr | Left Marker | dist (cM) | Right Marker | dist (cM) | Founder | Size | Founder Prob | Founder LOGP | Prob | % var | LOGP |
| --- | --- | --- | --- | --- | --- | --- | --- | --- | --- | --- | --- |
| 1B | IAAV5516 | 297.84 | BS00001128_51 | 298.34 | 1 | -0.207 | 0.399 | 0.4 | 0.048 | 1.6 | 1.32 |
|  |  |  |  |  | 2 | 0.946 | 0.077 | 1.11 |  |  |  |
|  |  |  |  |  | 3 | 0.112 | 0.445 | 0.35 |  |  |  |
|  |  |  |  |  | 4 | 0.318 | 0.304 | 0.52 |  |  |  |
|  |  |  |  |  | 5 | 0.698 | 0.148 | 0.83 |  |  |  |
|  |  |  |  |  | 6 | -1.119 | 0.07 | 1.15 |  |  |  |
|  |  |  |  |  | 7 | -1.185 | 0.021 | 1.67 |  |  |  |
|  |  |  |  |  | 8 | 0.437 | 0.259 | 0.59 |  |  |  |
| 1D | Kukri_c2464_592 | 18.04 | D_contig57829_235 | 18.21 | 1 | -0.81 | 0.113 | 0.95 | 0.003 | 2.5 | 2.59 |
|  |  |  |  |  | 2 | -0.569 | 0.238 | 0.62 |  |  |  |
|  |  |  |  |  | 3 | 1.282 | 0.042 | 1.37 |  |  |  |
|  |  |  |  |  | 4 | -1.024 | 0.066 | 1.18 |  |  |  |
|  |  |  |  |  | 5 | 0.631 | 0.27 | 0.57 |  |  |  |
|  |  |  |  |  | 6 | 0.849 | 0.12 | 0.92 |  |  |  |
|  |  |  |  |  | 7 | -1.18 | 0.036 | 1.45 |  |  |  |
|  |  |  |  |  | 8 | 0.82 | 0.21 | 0.68 |  |  |  |
| 1D | CAP11_c1157_168 | 100.79 | Kukri_c29687_369 | 100.95 | 1 | 0.552 | 0.254 | 0.6 | 0.275 | 0.9 | 0.56 |
|  |  |  |  |  | 2 | 0.105 | 0.446 | 0.35 |  |  |  |
|  |  |  |  |  | 3 | -0.11 | 0.447 | 0.35 |  |  |  |
|  |  |  |  |  | 4 | 0.021 | 0.49 | 0.31 |  |  |  |
|  |  |  |  |  | 5 | -0.044 | 0.477 | 0.32 |  |  |  |
|  |  |  |  |  | 6 | 0.003 | 0.498 | 0.3 |  |  |  |
|  |  |  |  |  | 7 | 0.734 | 0.134 | 0.87 |  |  |  |
|  |  |  |  |  | 8 | -1.261 | 0.015 | 1.82 |  |  |  |
| 2A | RAC875_c20273_823 | 35.38 | BS00094817_51 | 35.88 | 1 | 0.45 | 0.218 | 0.66 | 0.035 | 1.6 | 1.45 |
|  |  |  |  |  | 2 | 0.771 | 0.163 | 0.79 |  |  |  |
|  |  |  |  |  | 3 | 0.133 | 0.418 | 0.38 |  |  |  |
|  |  |  |  |  | 4 | -1.126 | 0.032 | 1.5 |  |  |  |
|  |  |  |  |  | 5 | -0.193 | 0.404 | 0.39 |  |  |  |
|  |  |  |  |  | 6 | -1.262 | 0.019 | 1.71 |  |  |  |
|  |  |  |  |  | 7 | 0.739 | 0.11 | 0.96 |  |  |  |
|  |  |  |  |  | 8 | 0.487 | 0.241 | 0.62 |  |  |  |
| 2B | BS00060618_51 | 317.61 | BS00000012_51 | 318.61 | 1 | 0.176 | 0.417 | 0.38 | 0.049 | 1.6 | 1.31 |
|  |  |  |  |  | 2 | 0.748 | 0.175 | 0.76 |  |  |  |
|  |  |  |  |  | 3 | 1.643 | 0.022 | 1.66 |  |  |  |
|  |  |  |  |  | 4 | -0.88 | 0.107 | 0.97 |  |  |  |
|  |  |  |  |  | 5 | -0.565 | 0.206 | 0.69 |  |  |  |
|  |  |  |  |  | 6 | -0.392 | 0.267 | 0.57 |  |  |  |
|  |  |  |  |  | 7 | -0.326 | 0.304 | 0.52 |  |  |  |
|  |  |  |  |  | 8 | -0.404 | 0.265 | 0.58 |  |  |  |
| 2D | Kukri_c27309_590 | 48.57 | BS00064538_51 | 56.64 | 1 | 0.669 | 0.293 | 0.53 | 0 | 18.5 | 18.33 |
|  |  |  |  |  | 2 | 2.727 | 0.009 | 2.03 |  |  |  |
|  |  |  |  |  | 3 | 0.215 | 0.44 | 0.36 |  |  |  |
|  |  |  |  |  | 4 | 1.112 | 0.211 | 0.68 |  |  |  |
|  |  |  |  |  | 5 | 0.759 | 0.276 | 0.56 |  |  |  |
|  |  |  |  |  | 6 | 0.897 | 0.245 | 0.61 |  |  |  |
|  |  |  |  |  | 7 | -6.194 | 0 | 7.11 |  |  |  |
|  |  |  |  |  | 8 | -0.185 | 0.437 | 0.36 |  |  |  |
| 3A | wsnp_Ex_rep_c106152_90334299 | 78.8 | Kukri_c51247_322 | 83.81 | 1 | 0.671 | 0.184 | 0.74 | 0.001 | 3 | 2.84 |
|  |  |  |  |  | 2 | -1.839 | 0.006 | 2.25 |  |  |  |
|  |  |  |  |  | 3 | -0.671 | 0.187 | 0.73 |  |  |  |
|  |  |  |  |  | 4 | 0.716 | 0.155 | 0.81 |  |  |  |
|  |  |  |  |  | 5 | -0.885 | 0.102 | 0.99 |  |  |  |
|  |  |  |  |  | 6 | 0.215 | 0.374 | 0.43 |  |  |  |
|  |  |  |  |  | 7 | 0.247 | 0.364 | 0.44 |  |  |  |
|  |  |  |  |  | 8 | 1.546 | 0.01 | 2.01 |  |  |  |
| 3D | RAC875_c48773_253 | 31.11 | RFL_Contig2471_119 | 31.28 | 1 | -0.133 | 0.442 | 0.35 | 0.417 | 0.7 | 0.38 |
|  |  |  |  |  | 2 | 0.159 | 0.431 | 0.37 |  |  |  |
|  |  |  |  |  | 3 | -0.351 | 0.351 | 0.45 |  |  |  |
|  |  |  |  |  | 4 | -0.525 | 0.276 | 0.56 |  |  |  |
|  |  |  |  |  | 5 | -0.025 | 0.489 | 0.31 |  |  |  |
|  |  |  |  |  | 6 | -0.443 | 0.244 | 0.61 |  |  |  |
|  |  |  |  |  | 7 | 1.191 | 0.021 | 1.68 |  |  |  |
|  |  |  |  |  | 8 | 0.128 | 0.423 | 0.37 |  |  |  |
| 4A | BS00064140_51 | 133.18 | BS00101512_51 | 134.18 | 1 | 0.521 | 0.273 | 0.56 | 0.002 | 2.4 | 2.75 |
|  |  |  |  |  | 2 | 0.176 | 0.415 | 0.38 |  |  |  |
|  |  |  |  |  | 3 | 0.277 | 0.376 | 0.42 |  |  |  |
|  |  |  |  |  | 4 | 0.266 | 0.36 | 0.44 |  |  |  |
|  |  |  |  |  | 5 | 0.118 | 0.447 | 0.35 |  |  |  |
|  |  |  |  |  | 6 | 0.019 | 0.488 | 0.31 |  |  |  |
|  |  |  |  |  | 7 | 0.727 | 0.187 | 0.73 |  |  |  |
|  |  |  |  |  | 8 | -2.104 | 0 | 3.63 |  |  |  |
| 5B | BobWhite_c22572_782 | 196.48 | Kukri_c836_513 | 196.65 | 1 | -3.398 | 0.001 | 3.11 | 0 | 6 | 3.85 |
|  |  |  |  |  | 2 | 0.725 | 0.232 | 0.63 |  |  |  |
|  |  |  |  |  | 3 | 0.886 | 0.158 | 0.8 |  |  |  |
|  |  |  |  |  | 4 | 1.621 | 0.088 | 1.05 |  |  |  |
|  |  |  |  |  | 5 | -0.001 | 0.5 | 0.3 |  |  |  |
|  |  |  |  |  | 6 | -1.505 | 0.037 | 1.43 |  |  |  |
|  |  |  |  |  | 7 | 0.704 | 0.206 | 0.69 |  |  |  |
|  |  |  |  |  | 8 | 0.968 | 0.141 | 0.85 |  |  |  |
| 6B | wsnp_Ex_c12450_19850827 | 215.75 | tplb0045b09_1555 | 216.42 | 1 | 0.089 | 0.461 | 0.34 | 0.269 | 1.1 | 0.57 |
|  |  |  |  |  | 2 | 0.723 | 0.117 | 0.93 |  |  |  |
|  |  |  |  |  | 3 | -0.552 | 0.264 | 0.58 |  |  |  |
|  |  |  |  |  | 4 | 0.368 | 0.333 | 0.48 |  |  |  |
|  |  |  |  |  | 5 | -0.111 | 0.447 | 0.35 |  |  |  |
|  |  |  |  |  | 6 | 0.247 | 0.388 | 0.41 |  |  |  |
|  |  |  |  |  | 7 | 0.467 | 0.299 | 0.53 |  |  |  |
|  |  |  |  |  | 8 | -1.232 | 0.02 | 1.71 |  |  |  |
| 6D | BS00094895_51 | 124.37 | GENE_4168_925 | 125.04 | 1 | -0.044 | 0.472 | 0.33 | 0.169 | 1 | 0.77 |
|  |  |  |  |  | 2 | -0.093 | 0.449 | 0.35 |  |  |  |
|  |  |  |  |  | 3 | 0.85 | 0.082 | 1.09 |  |  |  |
|  |  |  |  |  | 4 | -0.622 | 0.216 | 0.66 |  |  |  |
|  |  |  |  |  | 5 | -0.674 | 0.198 | 0.7 |  |  |  |
|  |  |  |  |  | 6 | 0.592 | 0.169 | 0.77 |  |  |  |
|  |  |  |  |  | 7 | -0.56 | 0.152 | 0.82 |  |  |  |
|  |  |  |  |  | 8 | 0.551 | 0.187 | 0.73 |  |  |  |
| 7B | BS00036788_51 | 58.06 | CAP7_c10566_170 | 58.22 | 1 | -0.73 | 0.18 | 0.74 | 0.008 | 2.5 | 2.09 |
|  |  |  |  |  | 2 | -0.247 | 0.402 | 0.4 |  |  |  |
|  |  |  |  |  | 3 | 0.422 | 0.312 | 0.51 |  |  |  |
|  |  |  |  |  | 4 | 1.264 | 0.029 | 1.53 |  |  |  |
|  |  |  |  |  | 5 | 0.22 | 0.411 | 0.39 |  |  |  |
|  |  |  |  |  | 6 | 1.249 | 0.065 | 1.19 |  |  |  |
|  |  |  |  |  | 7 | -0.704 | 0.173 | 0.76 |  |  |  |
|  |  |  |  |  | 8 | -1.473 | 0.012 | 1.91 |  |  |  |

**Supplemental Table S2:** **The chr1B_685645813/** **Ser674Gly SNP is globally rare.** Wheat HapMap data of a diverse germplasm collection and the 10+ Wheat genome project (italics). UK varieties highlighted in bold. Genotype of A|A corresponds to homozygous for adenine at the Ser674Gly SNP encoding the ancestral serine whilst G|G corresponds to homozygous guanine encoding the non-ancestral glycine (red).

| **Variety** | **Genotype (forward strand)** | **Variety** | **Genotype (forward strand)** | **Variety** | **Genotype (forward strand)** |
| --- | --- | --- | --- | --- | --- |
| 93 | A\|A | *Jagger* | *A\|A* | PI278297 | A\|A |
| 102 | A\|G | *Julius* | *A\|A* | PI349512 | A\|A |
| 403 | A\|A | klein_chamaco | A\|A | PI366716 | A\|A |
| 407-IV_60 | A\|A | *Lancer* | *A\|A* | PI366905 | A\|A |
| AC_Barrie | A\|A | M6 | A\|A | PI382150 | A\|A |
| acc1 | A\|A | *Mace* | *A\|A* | PI406517 | A\|A |
| acc2 | A\|A | Marquis | A\|A | PI445736 | A\|A |
| acc3 | A\|G | Neepawa | A\|A | PI470817 | A\|A |
| acc4 | A\|G | *Norin61* | *A\|A* | PI477870 | A\|A |
| acc5 | A\|A | Opata | A\|A | PI481718 | A\|A |
| Alabasskaja | A\|A | ***Paragon*** | ***G\|G*** | PI481923 | A\|A |
| *Arina* | *G\|G* | pavon | A\|A | PI565213 | A\|A |
| **Avalon** | **A\|A** | pbw343 | A\|A | PI82469 | A\|A |
| ***Cadenza*** | ***G\|G*** | PI153785 | A\|A | PI8813 | A\|A |
| *CDC Landmark* | *A\|A* | PI166180 | A\|A | PR267 | A\|A |
| *CDC Stanley* | *A\|A* | PI166333 | A\|A | rac875 | A\|A |
| chakwat86 | A\|A | PI177943 | A\|A | **Rialto** | **G\|G** |
| cham6 | A\|A | PI185715 | A\|A | ***Robigus*** | ***A\|A*** |
| Chinese Spring | A\|A | *PI190962* | *A\|A* | *Sy Mattis* | *A\|A* |
| ***Claire*** | ***A\|A*** | PI192001 | A\|A | **Taxi** | **A\|A** |
| clear_white | A\|A | PI192569 | A\|A | Truman | A\|A |
| dharwar_dry | A\|A | PI222669 | A\|A | Utmost | A\|A |
| Estacao | A\|A | PI245368 | A\|A | vorobey | A\|A |
| hidhab | A\|A | PI262611 | A\|A | *Weebil* | *A\|A* |

**Supplemental Table S3: The Ser674Gly SNP segregates with heading date in UK varieties.** Additional genotyping and heading information from Zikhali *et al*., 2016 (Zikhali et al., 2016) TGAC reference sequences (Clavijo et al., 2017). MAGIC founder varieties highlighted in bold, non-ancestral glycine in red.

| **Variety** | **Allele** | **Heading** |
| --- | --- | --- |
| **Alchemy** | A\|A | early |
| **Claire** | A\|A | early |
| **Robigus** | A\|A | early |
| Avalon | A\|A | early |
| **Brompton** | G\|G | late |
| **Hereward** | G\|G | late |
| **Rialto** | G\|G | late |
| **Xi-19** | G\|G | late |
| Paragon | G\|G | NA |
| Cadenza | G\|G | late |
| Spark | G\|G | NA |
| Badger | G\|G | NA |
| Savannah | G\|G | NA |

**Supplemental Table S4: Haplotype analysis of the *TaELF3* region between exon 2 and exon 3 (see Fig 3).** Variant location refers to position relative to Chinese spring *TaELF3-1D* transcription start site, first letter refers to base in Chinese Spring, second letter refers to changed base.

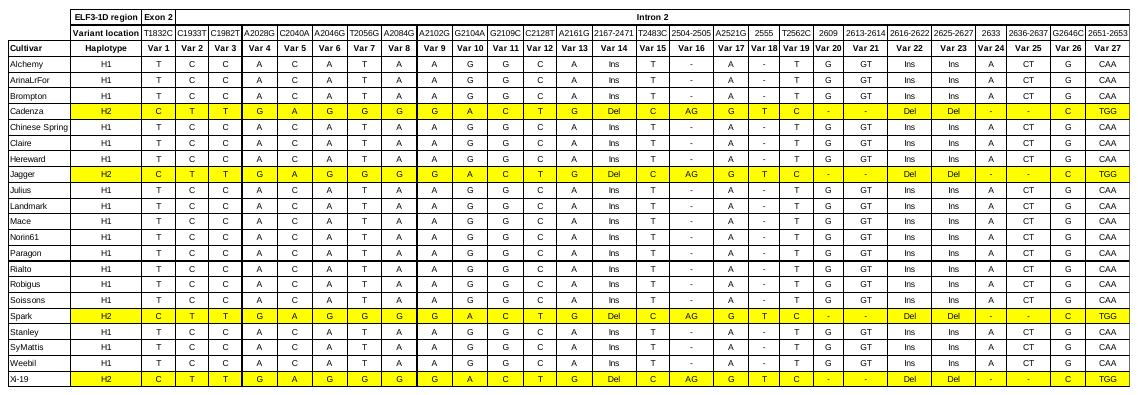

**Supplemental Table S5: Alignment of 300kb surrounding *ELF3-1D* in 10+ wheat genome lines.** Average nucleotide identity scores from alignment of 300kb sequence surrounding *ELF3-1D* in the 10+ wheat genome cultivars. Alignment produced using the whole genome alignment tool from CLC workbench 21.0.3 using a 15-base seed with no mismatches.

| **Cultivar** | **ArinaLrFor** | **CDC_Landmark** | **CDC_Stanley** | **Lancer** | **Mace** | **Norin61** | **CSv1.0** | **Julius** | **SYMattis** | **Jagger** | **Cadenza** |
| --- | --- | --- | --- | --- | --- | --- | --- | --- | --- | --- | --- |
| **ArinaLrFor** | 100.00 | 99.82 | 99.94 | 99.90 | 99.95 | 99.81 | 99.89 | 99.93 | 99.97 | 91.00 | 91.44 |
| **CDC_Landmark** | 99.82 | 100.00 | 100.00 | 99.89 | 99.85 | 99.93 | 99.82 | 99.90 | 99.94 | 90.57 | 91.44 |
| **CDC_Stanley** | 99.94 | 100.00 | 100.00 | 99.89 | 99.95 | 99.97 | 99.89 | 99.92 | 99.94 | 90.76 | 91.44 |
| **Lancer** | 99.90 | 99.89 | 99.89 | 100.00 | 99.94 | 99.92 | 99.91 | 99.94 | 99.93 | 91.00 | 91.42 |
| **Mace** | 99.95 | 99.85 | 99.95 | 99.94 | 100.00 | 99.90 | 99.86 | 99.98 | 99.98 | 90.51 | 91.44 |
| **Norin61** | 99.81 | 99.93 | 99.97 | 99.92 | 99.90 | 100.00 | 99.95 | 99.93 | 99.95 | 91.01 | 91.43 |
| **CSv1.0** | 99.89 | 99.82 | 99.89 | 99.91 | 99.86 | 99.95 | 100.00 | 99.93 | 99.83 | 91.00 | 91.44 |
| **Julius** | 99.93 | 99.90 | 99.92 | 99.94 | 99.98 | 99.93 | 99.93 | 100.00 | 99.99 | 91.00 | 91.43 |
| **SYMattis** | 99.97 | 99.94 | 99.94 | 99.93 | 99.98 | 99.95 | 99.83 | 99.99 | 100.00 | 91.00 | 91.43 |
| **Jagger** | 91.00 | 90.57 | 90.76 | 91.00 | 90.51 | 91.01 | 91.00 | 91.00 | 91.00 | 100.00 | 99.94 |
| **Cadenza** | 91.44 | 91.44 | 91.44 | 91.42 | 91.44 | 91.43 | 91.44 | 91.43 | 91.43 | 99.94 | 100.00 |

**Supplemental Table S6: Promoter motif analysis of *TtELF3.*** Column A refers to the A sub-genome, B sub-genome and D to the D sub-genome. Motifs and corresponding sequences taken from (Gendron et al., 2012; Huang et al., 2012; Wang et al., 1997)

| **Motif** | **Abbreviation** | **Sequence** | **A** | **B** | **D** | **Arabidopsis** |
| --- | --- | --- | --- | --- | --- | --- |
| Evening element | EE | AAAATATCT | 0 | 1 | 0 | 0 |
| Evening Like element | ELE | AATATCT | 1 | 0 | 0 | 1 |
| CCA1 binding site (A variant) | / | AAAAATCT | 0 | 0 | 0 | 0 |
| CCA1 binding site (C variant) | / | AACAATCT | 0 | 0 | 0 | 2 |
| Morning element 1 | ME 1 | CCACAC | 0 | 1 | 1 | 0 |
| Morning element 2 | ME 2 | GTGTGG | 1 | 1 | 2 | 0 |
| Hormone Up at Dawn 1 | HUD 1 | CACATG | 2 | 3 | 4 | 2 |
| Hormone Up at Dawn 2 | HUD 2 | CATGTG | 3 | 9 | 6 | 0 |
| G BOX (morning) | / | CACGTG | 1 | 0 | 3 | 0 |
| GATA (evening) | / | GGATA | 4 | 8 | 6 | 6 |
| Starch box (midnight) | / | AAGCCC | 1 | 0 | 1 | 0 |
| Telo-Box (midnight) | / | AAACCCT | 2 | 0 | 0 | 0 |
| Protein Box (midnight) | / | ATGGGCC | 0 | 2 | 1 | 0 |

**Supplemental Table S7: Promoter motif analysis of *TtLHY.*** Column A refers to the A sub-genome, B sub-genome and D to the D sub-genome. Motifs and corresponding sequences taken from (Gendron et al., 2012; Huang et al., 2012; Wang et al., 1997).

| **Motif** | **Abbreviation** | **Sequence** | **A** | **B** | **D** | **Arabidopsis** |
| --- | --- | --- | --- | --- | --- | --- |
| Evening element | EE | AAAATATCT | 0 | 0 | 0 | 0 |
| Evening Like element | ELE | AATATCT | 0 | 0 | 0 | 0 |
| CCA1 binding site (A variant) | / | AAAAATCT | 0 | 1 | 0 | 0 |
| CCA1 binding site (C variant) | / | AACAATCT | 1 | 0 | 0 | 0 |
| Morning element 1 | ME 1 | CCACAC | 1 | 1 | 1 | 1 |
| Morning element 2 | ME 2 | GTGTGG | 2 | 0 | 0 | 0 |
| Hormone Up at Dawn 1 | HUD 1 | CACATG | 1 | 1 | 1 | 1 |
| Hormone Up at Dawn 2 | HUD 2 | CATGTG | 4 | 3 | 3 | 0 |
| G BOX (morning) | / | CACGTG | 1 | 1 | 0 | 1 |
| GATA (evening) | / | GGATA | 3 | 3 | 3 | 3 |
| Starch box (midnight) | / | AAGCCC | 0 | 0 | 1 | 1 |
| Telo-Box (midnight) | / | AAACCCT | 0 | 0 | 0 | 0 |
| Protein Box (midnight) | / | ATGGGCC | 0 | 1 | 1 | 0 |

**Supplemental Table S8:**  **Promoter motif analysis of *TtLUX.*** Column A refers to the A sub-genome, B sub-genome and D to the D sub-genome. Motifs and corresponding sequences taken from (Gendron et al., 2012; Huang et al., 2012; Wang et al., 1997).

| **Motif** | **Abbreviation** | **Sequence** | **A** | **B** | **D** | **Arabidopsis** |
| --- | --- | --- | --- | --- | --- | --- |
| Evening element | EE | AAAATATCT | 9 | 4 | 4 | 1 |
| Evening Like element | ELE | AATATCT | 3 | 1 | 1 | 2 |
| CCA1 binding site (A variant) | / | AAAAATCT | 1 | 0 | 0 | 2 |
| CCA1 binding site (C vairiant) | / | AACAATCT | 0 | 1 | 0 | 0 |
| Morning element 1 | ME 1 | CCACAC | 1 | 1 | 1 | 0 |
| Morning element 2 | ME 2 | GTGTGG | 1 | 1 | 0 | 0 |
| Hormone Up at Dawn 1 | HUD 1 | CACATG | 2 | 1 | 2 | 0 |
| Hormone Up at Dawn 2 | HUD 2 | CATGTG | 2 | 2 | 0 | 1 |
| G BOX (morning) | / | CACGTG | 2 | 3 | 2 | 2 |
| GATA (evening) | / | GGATA | 3 | 5 | 2 | 2 |
| Starch box (midnight) | / | AAGCCC | 0 | 0 | 2 | 0 |
| Telo-Box (midnight) | / | AAACCCT | 3 | 2 | 1 | 2 |
| Protein Box (midnight) | / | ATGGGCC | 1 | 1 | 2 | 0 |

**Supplemental Table S9: Total number of SNPs within the coding sequence of ELF3 in the 10+ Wheat Genomes project cultivars.** A SNP was recorded if the base was different to the Chinese Spring IWGSCRefseqv1.1 sequence in the alignment. SNPs within the same codon were recorded individually. It was not possible to create clear alignments between ELF3-1D Chinese Spring, Jagger and Cadenza and so the coding sequence for the three cultivars was downloaded from ENSEMBL plants, aligned and the number of SNPs manually counted. NS = non-synonymous SNP, S= synonymous SNP, PAV = presence/absence variation. Cultivars were considered to share the same allele if the CDS was identical. Zavitan is a tetraploid variety and so lacks a D sub genome.

|  | **Chromosome** | | | | | |
| --- | --- | --- | --- | --- | --- | --- |
|  | **1A** | | **1B** | | **1D** | |
| **Cultivar** | **NS** | **S** | **NS** | **S** | **NS** | **S** |
| ArinaLrFor | 2 | 2 | 1 | 0 | 0 | 0 |
| CDC Stanley | 2 | 2 | 0 | 0 | 0 | 0 |
| CDC Landmark | 2 | 2 | 0 | 0 | 0 | 0 |
| Cadenza | 0 | 1 | 1 | 0 | 35 | 23 |
| Claire | 0 | 1 | 0 | 0 | 0 | 0 |
| Jagger | 2 | 3 | 0 | 1 | 35 | 23 |
| Julius | PAV | PAV | 0 | 0 | 1 | 0 |
| Lancer | 2 | 2 | 0 | 0 | 0 | 0 |
| Norin61 | 2 | 2 | 0 | 0 | 0 | 0 |
| Mace | 2 | 2 | 0 | 0 | 0 | 0 |
| PI190962 | 0 | 0 | 0 | 0 | 0 | 0 |
| Paragon | 0 | 1 | 1 | 0 | 0 | 0 |
| Robigus | 0 | 0 | 0 | 0 | 0 | 0 |
| SY Mattis | 0 | 1 | 0 | 0 | 0 | 0 |
| Zavitan | 7 | 2 | 1 | 3 | *NA* | *NA* |
| **Total No. SNP's** | 21 | 21 | 3 | 4 | 71 | 46 |
| **Total No. Alleles** | 5 | | 3 | | 3 | |
